# The calcium binding protein S100β marks hedgehog-responsive resident vascular stem cells within vascular lesions

**DOI:** 10.1101/2020.05.20.105981

**Authors:** Mariana Di Luca, Emma Fitzpatrick, Denise Burtenshaw, Weimin Liu, Jay-Christian Helt, Roya Hakimjavadi, Eoin Corcoran, Yusof Gusti, Daniel Sheridan, Susan Harman, Catriona Lally, Eileen M. Redmond, Paul A. Cahill

## Abstract

A hallmark of subclinical atherosclerosis is the accumulation of vascular smooth muscle cell (SMC)-like cells leading to intimal thickening. While medial SMCs contribute, the participation of hedgehog responsive resident vascular stem cells (vSCs) to lesion formation remains unclear. Using transgenic eGFP mice and genetic lineage tracing of S100β vSCs *in vivo*, we identified S100β/Sca1 cells derived from a S100β non-SMC parent population within lesions that co-localise with smooth muscle ⍰-actin (SMA) cells following iatrogenic flow restriction, an effect attenuated following hedgehog inhibition with the smoothened inhibitor, cyclopamine. *In vitro*, S100β/Sca1 cells isolated from atheroprone regions of the mouse aorta expressed hedgehog signalling components, acquired the di-methylation of histone 3 lysine 4 (H3K4me2) stable SMC epigenetic mark at the Myh11 locus and underwent myogenic differentiation in response to recombinant sonic hedgehog (SHh). Both S100β and PTCH1 cells were present in human vessels while S100β cells were enriched in arteriosclerotic lesions. Recombinant SHh promoted myogenic differentiation of human induced pluripotent stem cell-derived S100β neuroectoderm progenitors *In vitro*. We conclude that hedgehog responsive S100β vSCs contribute to lesion formation and support targeting hedgehog signalling to treat subclinical arteriosclerosis.

## Introduction

Atherosclerosis is a chronic progressive inflammatory disease that leads to myocardial infarction and stroke ^1^. Pathologic observations in humans vessels confirm that early ‘transitional’ lesions enriched with smooth muscle (SMC)-like cells called adaptive lesions are routinely present in atherosclerotic-prone regions of arteries before pathologic intimal thickening, lipid retention, and the appearance of a developed plaque ^2^. Endothelial cell (EC) dysfunction due to disturbed blood flow is classically associated with the development of atheroprone lesions^3^, while the embryological origin (e.g., neuroectoderm versus mesoderm) of SMCs may also influence disease localisation and progression^4^. These early lesions are routinely modelled in mouse carotid arteries following flow restriction^5^ and can further rapidly develop into advanced atherosclerotic plaques in ApoE knockout on a Western diet ^6,7^.

Lineage tracing and single cell RNA sequence analysis (scRNA-seq) have provided compelling evidence in various animal models for the involvement of a rare population of (i) differentiated myosin heavy chain 11 (Myh11) medial SMC that are stem cell antigen-1 positive (Sca1^+^) ^8,9^ and (ii) various adventitial and medial progenitors in progressing intimal thickening ^10–13^. While some adventitial progenitors are derived from a parent Myh11 SMC population ^14^, non-SMC derived SRY-related HMG-box 10 (Sox10) adventitial/medial resident multipotent vascular stem cells (vSC), glioma-associated oncogene homolog 1 (Gli) and Nestin progenitor cells have also been implicated ^13,15^. Despite these insights, the putative role of resident vSCs that do not originate from medial Myh11^+^ SMCs in promoting intimal thickening remains controversial.

S100β, a member of the S100 multigenic family expressing small (9⍰Da and 14⍰Da) Ca2+- binding proteins of the EF-hand type, is primarily localised to astrocytes and glial cells under normal physiological conditions and is positively associated with age and various neurodegenerative and neuroinflammatory diseases ^16^. S100β promotes intimal thickening through interaction with the receptor for advanced glycation end-products (RAGE) ^17^ and induces migration and infiltration of inflammatory cells ^18,19^. S100β plays a crucial role in maintaining an intermediate state of Sca1 progenitor cells following injury-induced neointimal thickening through a RAGE and stromal cell-derived factor-1 α/CXCR4 signalling mechanism^20^.

Hedgehog signalling (Hh) is essential for normal embryonic development and plays a pivotal role in the maintenance of adult progenitor/stem cells in tissue repair after injury ^21^. Hh signalling is initiated upon binding of a ligand to the canonical receptors, PTCH1 and PTCH2 (Patched 1 and 2) ^22^, which leads to disinhibition of SMO (Smoothened) and activation of a complex signalling cascade that regulates the GLI (glioma-associated oncogene homolog 1) family zinc finger transcription factors (GLI1, GLI2, and GLI3) ^23^. GLI activators induce the transcription of Hh target genes primarily involved in cell proliferation, cell survival, and cell fate specification, including PTCH and GLI ^24,25^. HHIP (hedgehog interacting protein) modulates hedgehog signalling activity by binding and inhibiting the action of hedgehog proteins ^26^. Genome wide association studies (GWAS) in coronary artery disease patients have identified variants at the 14q32 HHIPL1 (Hedgehog interacting protein-like 1) that encode a paralog of HHIP ^27^ to increase Hh signalling and regulate SMC proliferation and migration *In vitro* ^28^. In atherosclerotic mice, Hh has a permissive role through increased lipid uptake by macrophages ^29^, while global Hhipl1 deletion reduces plaque burden ^28^. Hh signalling, especially SHh and signal peptide CUB domain and EGF like domain containing 2 (Scube2), are overexpressed in injured arteries during lesion formation after flow restriction ^30–32^, autogenous vein grafts ^33^, and hypoxic pulmonary arterial SMCs ^34^. Adventitial Sca1^+^ cells that co-localise with SHh and Ptch1 ^35^ significantly contribute to intimal thickening *in vivo* ^31,36,37^ while Hh inhibition with cyclopamine or local perivascular depletion of Ptch1 attenuates intimal thickening following iatrogenic flow-restriction *in vivo* ^38,39^.

Given the pivotal role of Hh in the maintenance of adult progenitor/stem cells, tissue repair and atherosclerosis ^21,28^, we sought to determine the role of Hh responsive S100β/Sca1 vSCs and their myogenic progeny in contributing to intimal thickening and the progression of adaptive lesions *in vivo* and, furthermore, whether Hh signalling directs murine S100β vSCs and hiPSC-derived S100β neuroectoderm stem cells to undergo proliferation and myogenic differentiation *In vitro*.

## Results

### Perivascular Sca1 and S100β stem vSCs are present in normal adult vessels

The presence of perivascular Sca1 and S100β vSCs was first confirmed in whole mounted vessels from Sca1-eGFP and S100β-Cre-tdT mice. A discrete number of Sca1-eGFP and S100β-tdT cells were present on collagen fibres within the adventitial layer of the thoracic aorta, when compared to wild-type (WT) controls [Suppl Figure 1a-d]. The cells were located predominantly at the aortic arch and at bifurcations with intercostal vessels, and their presence diminished along the descending and abdominal aorta [Suppl Figure 1b, c]. When the media was removed, Sca1-eGFP cells were clearly present in adventitial layer of the thoracic aorta [Suppl Figure 1d] and in perivascular regions of the carotid artery [Suppl Figure 1d].

### Sca1 and S100β cells are present in intimal lesions from murine carotid arteries following iatrogenic flow restriction

The contribution of Sca1 cells to intimal medial thickening following iatrogenic flow restriction was first assessed in Sca1-eGFP transgenic mice following partial ligation of the left carotid artery (LCA). Vessels were harvested on day 3, 7 and 14 days post-ligation and paraformaldehyde fixed and paraffin-embedded cross sections were evaluated for eGFP (indicative of Sca1 expression) by confocal microscopy. Morphometric analysis confirmed that adventitial and intimal volumes increased significantly over time following ligation, without any significant change to medial volume, concomitant with a significant reduction in luminal volume after 7 and 14 days [Figure 1a].

**Figure 1.**
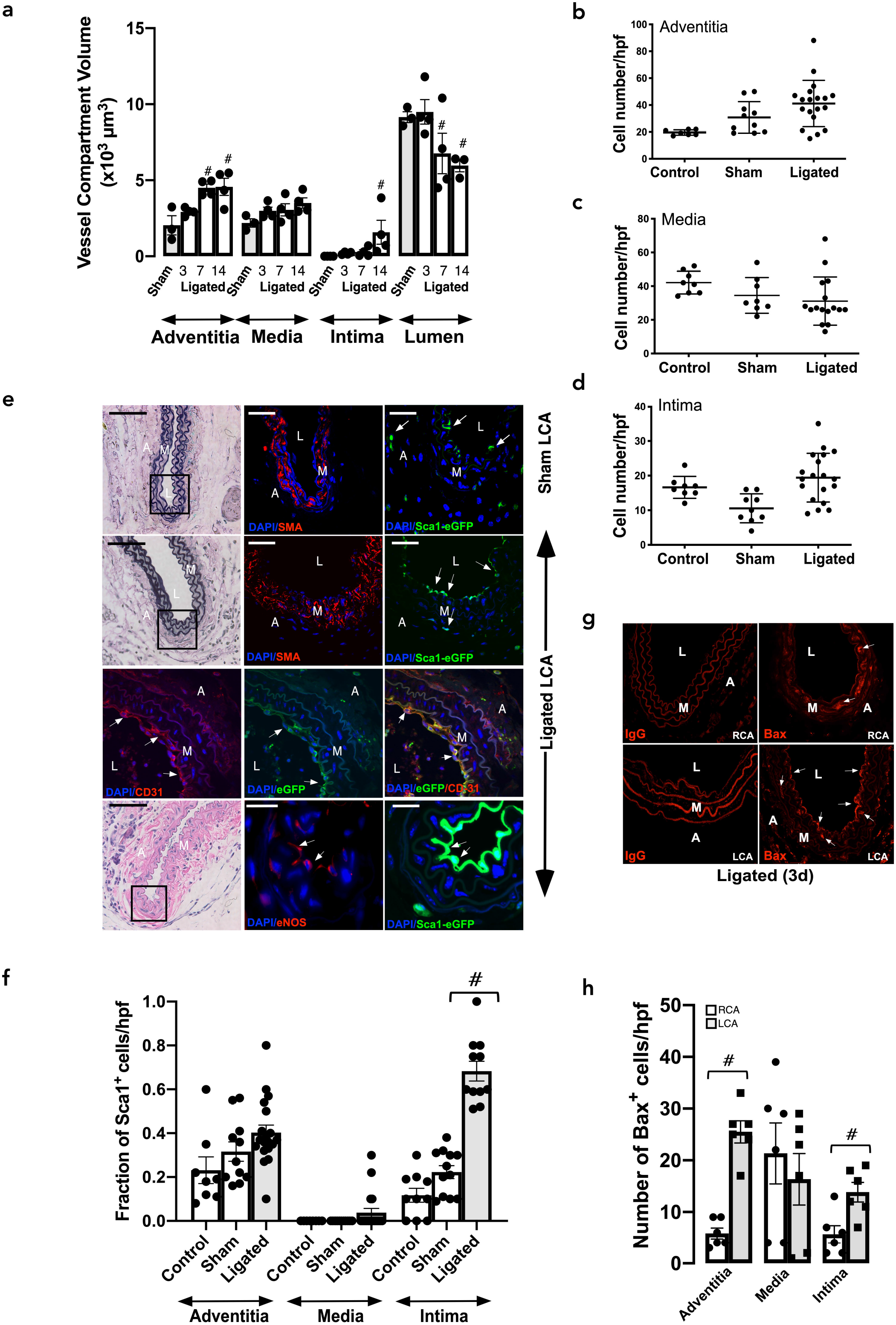
Expression of Sca1-eGFP and S100β-eGFP cells following iatrogenic flow restriction after 3 days. **a.** Morphometric analysis of adventitial, medial, intimal and luminal volumes following partial ligation of the left carotid artery compared to sham controls. Data are the mean ± SEM of 4 animals, #p<0.05 vs sham. **b-d.** The number of DAPI nuclei/hpf in cross-sections of (b) adventitia, (c) media and (d) intima following partial ligation of the left carotid artery (LCA) after 3d compared to untouched (control) and sham. Data are the mean ± SEM of 3-5 representative images per animals, n=3, #p<0.05. **e.** Verhoeff-Van Gieson stained sections and corresponding confocal fluorescence images of DAPI nuclei (blue), Sca1-eGFP (green) and immunofluorescence of IZ-actin (SMA) (red), anti-eGFP (green), anti-CD31 (red) and anti-eNOS (red) expression in ligated LCA vessels expression in sham-operated and ligated LCA vessels. Arrows highlight Sca1-eGFP expressing cells in the adventitia (A), media (M) and neointima (NI) and CD31 and eNOS in intima. Scale bar = 20μm. **f.** The fraction of Sca1 cells/high power field (hpf) in the adventitia, media, and intima of sham and ligated vessels after 3 days. Data are the mean ± SEM of 3-5 sections, #p≤ 0.05 from a minimum of 4 animals per experimental group. **g.** Representative immunofluorescence images for the pro-apoptotic marker, Bax, (red) in ligated left carotid artery (LCA) vessels and contralateral right carotid artery (RCA) after 3d. IgG was used as a control. **h.** The number of Bax cells/hpf in the adventitia, media, and intima of sham and ligated vessels after 3d. Arrows highlight Bax expressing cells within the adventitia (A), media (M), and neointima (NI). Data are the mean ± SEM of 3-6 sections per experimental group from 3 animals. #p≤ 0.05. Scale bar = 20μm.

The number of cells within the adventitial, medial and intimal layers of the LCA did not significantly change 3d post ligation [Figure 1b-d]. No Sca1-eGFP cells were visible within the medial layer of sham vessels which was populated primarily by ⍰-smooth muscle actin (SMA) SMCs [Figure 1e]. While a small but significant number of Sca1-eGFP cells were present within the adventitial and intimal layers of sham vessels, the fraction of Sca1-eGFP cells significantly increased within the intimal layer 3d post ligation [Figure 1e, f]. Immunohistochemical analysis using anti-eGFP confirmed the presence of intimal Sca1-eGFP cells in ligated vessels [Figure 1e] and further demonstrated that intimal Sca1-GFP cells co-express PECAM-1/CD31 and endothelial cell nitric oxide (eNOS) [Figure 1e]. Moreover, there was also a notable increase in the expression of the pro-apoptotic protein, Bax within the intimal, adventitial and medial layer of the LCA and the medial layer of the RCA 3d post ligation [Figure 1g,h], consistent with disturbed flow- and strain -induced apoptosis ^40^.

While the majority of Sca1-eGFP cells were present in the adventitial and intimal layers of sham operated vessels, a small fraction were also visible within the medial layer after 7, 14 and 21d [Figure 2a, b and g and Suppl Figure 2a, b]. Upon injury, there was a notable appearance of Sca1-eGFP cells within the expanding intimal layer of ligated vessels after 7, 14 and 21d [Figure 2a, b, d, g] that coincided with an increase in the number of cells [Figure 2c] and the volume of the neointimal layer [Figure 1a]. Cumulative analysis revealed that there was a significant increase in the fraction of Sca1-eGFP cells within neointimal layers after 7 and 14 and 21d, respectively, when compared to sham vessels or contralateral RCA [Figure 2d, h]. The majority of Sca1 cells within these lesions co-localised with SMA^+^ cells [Figure 2e, f, Suppl Figure 3a].

**Figure 2.**
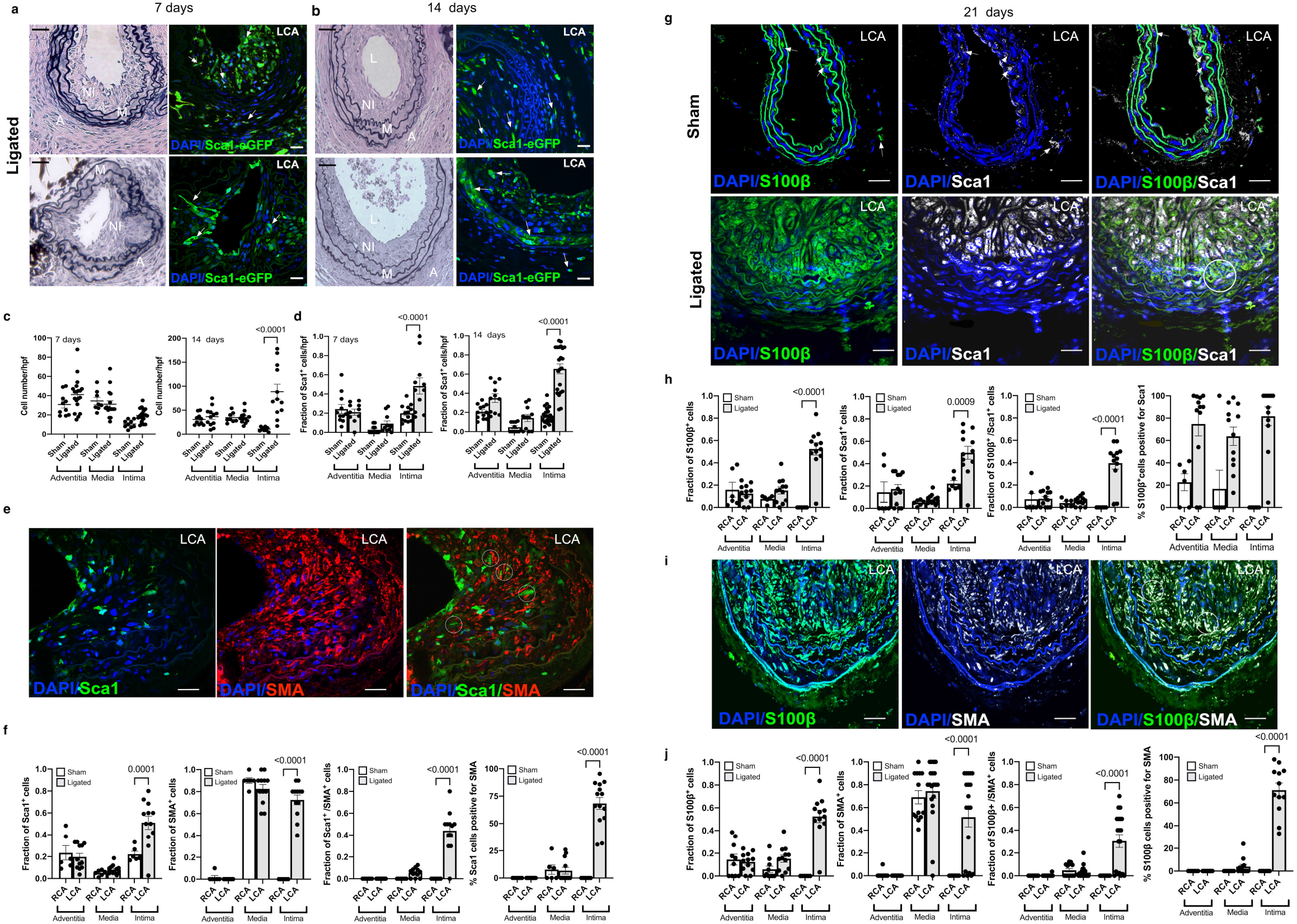
Expression of Sca1-eGFP and S100β-eGFP cells following iatrogenic flow restriction after 7, 14 and 21 days. **a,b.** Representative Verhoeff-Van Gieson stained sections and corresponding confocal fluorescence images of DAPI nuclei (blue) and Sca1-eGFP (green) expression in LCA after (a) 7 days and (b) 14 days. Scale bar = 20μm. **c.** The number of DAPI nuclei/hpf in cross-sections of adventitia, media and intima following partial ligation of the LCA after 7 and 14 days compared to sham controls. Data are the mean ± SEM of 3-5 representative images per animal, n=4. **d.** The fraction of Sca1 cells/high power field (hpf) in the adventitia, media, and intima of sham and ligated vessels after 7 days and 14 days. Data are the mean ± SEM of 10-15 sections from 4 animals per experimental group. **e.** Representative confocal fluorescence images of DAPI nuclei (blue), immunofluorescence of anti-eGFP (green) expression and anti-SMA (Red) in the LCA 14 days post-ligation in Sca1-eGFP mice. Data are representative of 5 images per animal. **f.** Cumulative analysis of Sca1^+^ cells, SMA^+^ cells, double stained Sca1/SMA positive cells and the percentage of Sca1 cells that were SMA1 in the adventitial, medial and intimal layer of RCA and LCA vessels after 21 days, respectively. Data are the mean ± SEM, n=15-20 sections from 5 animals. **g.** Representative confocal fluorescence images of DAPI nuclei (blue), immunofluorescence of anti-eGFP (green) expression and anti-Sca1 (far-red white) in sham and ligated LCA after 21 days in S100β-eGFP mice. Scale bar = 20μm. **h.** Cumulative analysis of S100β cells, Sca1 cells, double stained Sca1/S100β positive cells and the percentage of S100β cells that were Sca1 in the adventitial, medial and intimal layer of RCA and LCA vessels, respectively after 21 days in S100β-eGFP mice. Data are the mean ± SEM, n=15-20 sections from 5 animals/group. **i.** Representative confocal fluorescence images of DAPI nuclei (blue), immunofluorescence of anti-eGFP (green) expression and anti-SMA (far red, white) in the RCA and LCA 21 days post-ligation in S100β-eGFP mice. Data are from a minimum of 4 animals per experimental group. Scale bar = 20μm. **j.** Cumulative analysis of S100β cells, SMA cells, double stained S100β/SMA positive cells and the percentage of S100β cells that were SMA in the adventitial, medial and intimal layer of RCA and LCA vessels, respectively after 21 days in S100β-eGFP mice. Data are the mean ± SEM, n=15-20 sections from 4 animals/group.

The contribution of S100β vSC to intimal thickening following iatrogenic flow restriction was also assessed in S100β-eGFP transgenic mice. While the majority of S100β-eGFP cells were present in the adventitial layer of sham operated vessels and the contralateral RCA, a small fraction were also present within the medial layer after 21d [Suppl Figure 2a, c, Figure 2g, h]. In contrast, no intimal S100β-eGFP cells were present in sham vessels or the contralateral RCA [Suppl Figure 2a c-e]. Upon injury, the fraction of S100β-eGFP cells within lesions significantly increased when compared to sham vessels and the contralateral RCA [Suppl Figure 2d, e and Figure 2 g,h]. Importantly, the number of double positive Sca1/S100β cells increased within the neointimal layers of ligated vessels compared to the sham [Suppl Figure 2d, e] and contralateral RCA control [Figure 2g, h]. Similarly, the number of double positive S100β/SMA cells increased within lesions compared to the contralateral RCA [Figure 2i, j]. Taken together, these data suggest double positive Sca1/S100β cells populate vascular lesions following iatrogenic injury *in vivo*.

### Lineage tracing analysis of marked perivascular S100β cells iatrogenic flow restriction

In order to determine the source of intimal double positive Sca1/S100β cells following flow restriction, lineage tracing analysis was performed using transgenic S100β-eGFP-CreERT2-Rosa-26-tdTomato reporter mice [Suppl Figure 4a]. As cre-mediated recombination results in a permanent heritable change in a cell’s genome, these cells and their progeny remain tdTomato (tdT) positive. The animals were treated with tamoxifen (Tm) for 7 days to induce nuclear translocation of CreER(Tm) and subsequent recombination to indelibly mark S100β cells with red fluorescent tdT seven days before ligation was performed [Suppl Figure 4b]. Only S100β cells present during the period of Tm treatment and their subsequent progeny following injury are marked with tdT. The tissue specificity and recombination efficiency of the Tm-induced Cre activity was confirmed in bone-marrow smears and neuronal tissue from S100β-eGFP-CreERT2-Rosa-26-tdTomato mice. Treatment with Tm resulted in Cre-mediated recombination and expression of S100β-tdT in >90% of neuronal cells of the carotid sinus nerve (Hering’s nerve) without any S100β-tdT cells observed in the bone marrow smears [Suppl Figure 4c]. The neuronal staining was confirmed by immunohistochemical analysis of S100β expression using both anti-S100β and anti-tdT antibodies (data not shown).

Morphometric analysis of the S100β-eGFP-creER2-Rosa26-tdT mice confirmed significant intimal thickening in transgenic mice following ligation injury when compared to the sham LCA and contralateral RCA [Figure 3a-c]. Treatment of S100β-eGFP-creER2-Rosa26-tdT transgenic mice with tamoxifen indelibly marked S100β cells (S100β-tdT) within the adventitial layer of the sham LCA prior to flow restriction [Figure 3d]. No cells are marked when these double transgenic mice were treated with the vehicle control (corn oil) [Figure 3d]. Importantly, no S100β-tdT cells were observed in the intimal (EC) or medial (SMC) layers of these sham vessels following tamoxifen treatment [Figure 3d]. However, following ligation, there was a striking increase in the number of S100β-tdT marked cells within the medial and intimal layers of the ligated LCA, compared to sham controls [Figure 3e, f]. In order to rule out the possibility that Tm treatment continues to label a significant number of cells for weeks after last Tm treatment ^41^, we confirmed that the fraction of S100β-tdT marked cells remained significantly higher in the neointima of LCA four weeks after the last Tm injection when compared to the RCA [Suppl Figure 4d, e]. Moreover, the fraction of S100β-tdT marked cells within the LCA adventitial, medial and intimal layers of ligated vessels after one and four weeks from the last Tm injection was of a similar magnitude [Suppl Figure 4f].

**Figure 3.**
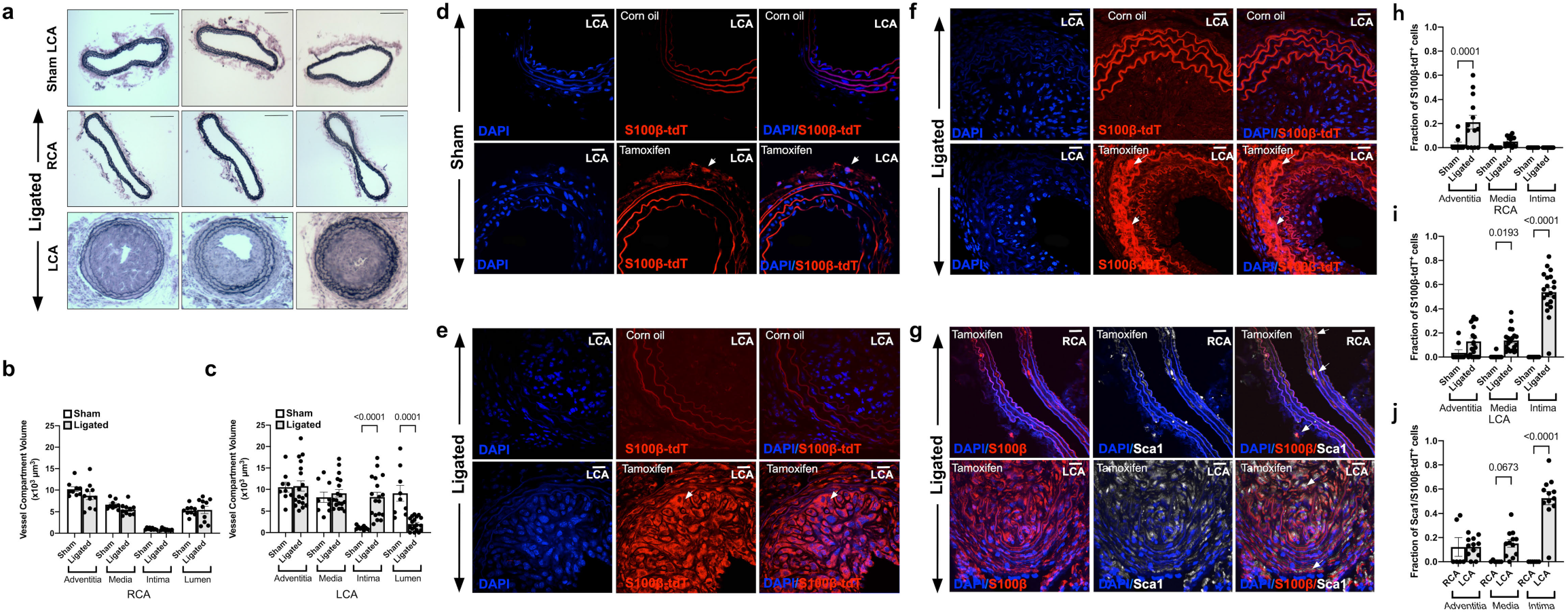
Lineage tracing analysis of S100β-Cre(ERT2)-tdT marked cells following iatrogenic flow restriction for 21 days. **a-c.** Verhoeff-Van Gieson stained sections and morphometric analysis of adventitial, medial, intimal and luminal volumes of the **(b)** contralateral RCA control compared to the **(c)** LCA following ligation for 21d. Data are the mean ± SEM of 10 images, #p<0.05 vs sham controls. **d-f.** Representative confocal fluorescence images of DAPI nuclei (blue) and S100β-tdT (red) cells in mice treated with corn oil (control) or tamoxifen for 7 days prior to washout for 1 week in **(d)** sham and **(e, f)** ligated LCA. Data are minimum of 4 animals per experimental group. Scale bar = 20 μm. Arrows highlight S100β-tdT (red) marked cells. **g.** Representative confocal fluorescence images of DAPI nuclei (blue), immunofluorescence of anti-tdT (red) and anti-Sca1 (far red, white) expression in the contralateral RCA and LCA 21 days post ligation. Scale bar = 20μm. **h-j.** Cumulative analysis of the fraction of S100β-tdT^+^ cells in the **(h)** RCA and **(i)** LCA and **(j)** Sca1/S100β double stained cells in the adventitial, medial and intimal layer of RCA and LCA vessels, respectively after 21 days in S100β-CreERT2-tdT mice.

When S100β-tdT marked cells were evaluated for co-expression of Sca1 and SMA by immunohistochemistry using anti-Sca1, anti-SMA and anti-tdT antibodies, S100β-tdT marked cells co-localised with Sca1 cells in the adventitial and medial layers (but not the intimal layer) of the control contralateral RCA before injury [Suppl Figure 3d, Figure 3g]. There was a notable increase in the number of double stained Sca1/S100β-tdT cells in the media and neointima of the LCA compared to the contralateral RCA [Figure 3g] and the sham LCA control [Suppl Figure 3b] in double stained SMA/S100β-tdT cells within the neointima and media compared to the sham control [Suppl Figure 3c]. Cumulative analysis confirmed both a significant increase in the fraction of S100β-tdT cells [Figure 3h] and the fraction of double stained S100β/Sca1 cells [Figure 3i] within the LCA medial and intimal layers compared to the contralateral RCA control. Moreover, the majority of Sca1 cells within ligated vessels were S100β-tdT cells [Figure 3j]. Since intimal and medial cells are not marked with tdT prior to injury, these data suggest that these cells are derived from a non-SMC S100β parent population following flow restriction. Moreover, the majority of intimal Sca1 cells are derived from an S100β parent population.

### Hedgehog signalling components control intimal thickening following iatrogenic flow restriction

The effect of systemic administration of the Hedgehog inhibitor cyclopamine (10 mg/kg, IP, every other day) on neointimal thickening and the accumulation of Sca1-eGFP cells was evaluated. 2-hydroxypropyl-β-cyclodextrin (HβCD) was used as a vehicle control. Vessels were harvested 7 and 14 d post-ligation and morphometric analysis was performed. Whole vessel Gli1 and Gli2 mRNA levels were significantly elevated in ligated vessels treated with HβCD when compared to sham controls (Gli1: 6.1 ± 1.3 after 3 d; Gli2:8.1 ± 0.9 fold after 14d, p≤ 0.05 n=4), an effect that was significantly reduced following cyclopamine treatment (Gli2: 3.2 ± 0.3 fold, n=4, p≤ 0.05). Immunohistochemical analysis revealed that the Hh receptor, Ptch1, was up-regulated in LCA following injury concomitant with enhanced Gli2 expression in LCA when compared to sham vessels [Figure 4a]. Gli2 co-localised with Sca1-eGFP expressing cells and was markedly increased in ligated vessels treated with vehicle (HβCD) when compared to sham-operated vessels, especially in the neointima and media of the LCA, an effect that was attenuated following Hh inhibition [Figure 4a]. Cyclopamine treatment significantly reduced the fraction of Sca1-eGFP expressing cells [Figure 4b, c] concomitant with reduced intimal thickening 7 d and 14 d-post ligation [Figure 4d-g]. Collectively, these data suggest that intimal thickening is attenuated by systemic administration of the smoothened inhibitor, cyclopamine, concomitant with a reduction in the accumulation of Gli2/Sca1 Hh-responsive cells within the intimal and medial layers.

**Figure 4.**
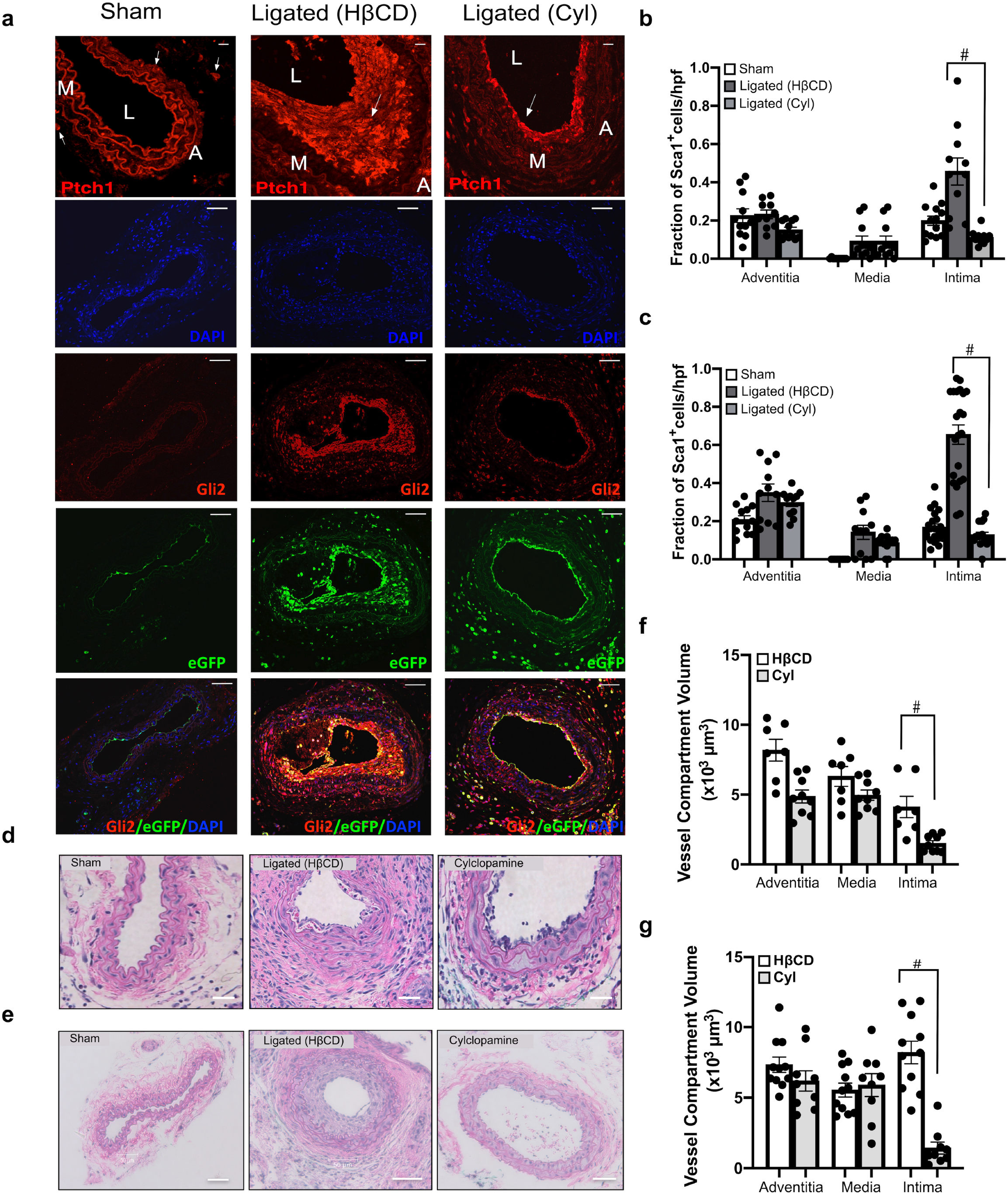
Inhibition of Hh signalling attenuates intimal thickening and Sca1 cell expansion following iatrogenic flow restriction. **a.** Representative immunofluorescence images of Hh receptor patched 1 (Ptch1) (red), Hh target gene Gli2 (red), Sca1-eGFP (green) expression and merged (eGFP/Gli2/DAPI) in carotid artery cross sections from sham-operated, ligated vehicle control (HβCD), and ligated + cyclopamine (10 mg/kg, IP) treated mice after 14 days. Scale bar = top panel: 50μm, all other panels: 20μm. **b, c.** Cumulative data showing the fraction of Sca1-eGFP cells within the adventitia, media and intima from sham-operated, ligated vehicle control (HβCD), and ligated + cyclopamine (Cyl) Sca1-eGFP mice after **(b)** 7 days and **(c)** 14 days. Data are the mean ± SEM of five sections per experimental group, #p≤ 0.05 vs ligated HβCD controls and are representative of a minimum of 4 animals per experimental group. **d, e.** Representative H&E staining of carotid artery cross sections from sham, ligated control (HβCD vehicle), and ligated + cyclopamine (Cyl) treated animals after 7 d **(d)** and **(e)** 14 d. Scale bar = 50 μm. **f,g.** Morphometric analysis of vessel compartment volumes for ligated control (HβCD), and ligated + cyclopamine (Cyl) treated animals after **(f)** 7d and **(g)** 14 d. Data mean ± SEM of 7-12 sections analysed from 4 animals, #p≤ 0.05 vs ligated control (HβCD).

### Isolation and characterisation of resident S100β/Sca1 vascular stem cells from murine aorta

The expression of SMC differentiation markers genes, *Myh11* and *Cnn1* was enriched in medial segments from both atheroprone aortic arch (AA) and atheroresistant thoracic/abdominal aorta (TA) regions of the mouse aorta, when compared to NE-4C cells and immortalised MOVAS mSMCs in culture [Figure 5a]. The level of S100β and Sox10 expression, both markers of resident vascular stem cells ^12,42^, was then assessed in AA and TA regions, with or without the adventitia [Figure 5b, c]. There was significant enrichment of both S100β and Sox10 in the AA when compared to the TA. Moreover, the levels were significantly enriched within the adventitia since its removal resulted in a dramatic reduction in the expression of both transcripts. Nevertheless, medial expression of both genes was still observed in the AA region [Figure 5b, c].

**Figure 5.**
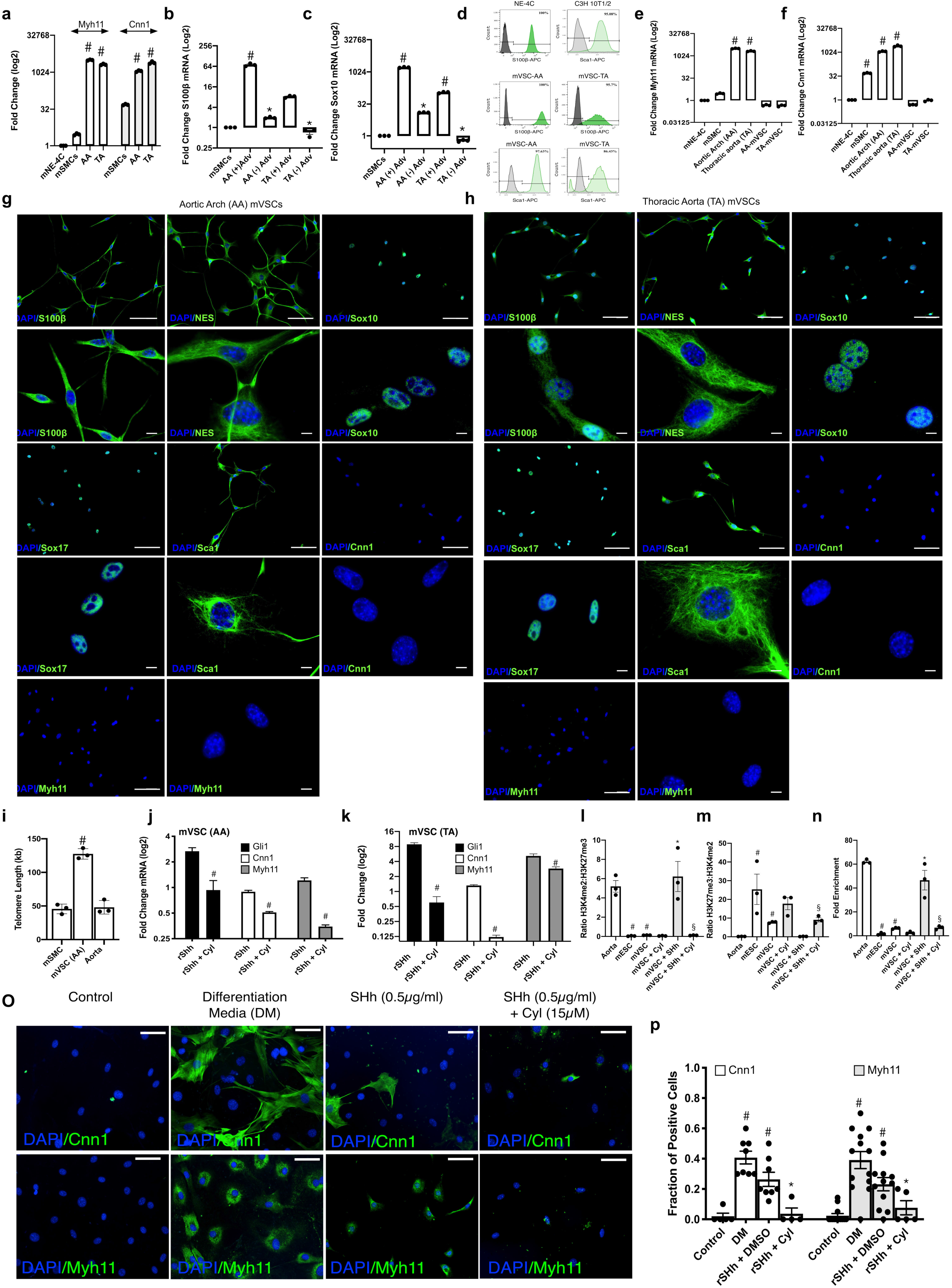
Resident S100β/Sca1 vascular stem cells from atheroprone and atheroresistant regions of the mouse aorta *In vitro*. **a.** Relative levels of *Myh11* and *Cnn1* in AA and TA regions of the mouse aorta. Data are expressed as the Log2 fold change in mRNA levels relative to neural stem cells (NE-4C) in culture and are the mean ± SEM of three aortic specimens, #p≤ 0.05 vs NE-4C cells. The housekeeping gene Gapdh was used as a control. **b, c.** The level of enrichment of mRNA for neuroectodermal markers **(b)** S100β and **(c)** Sox10 within atheroprone AA (aortic arch) and atheroresistant TA (thoracic/descending aorta) regions of the mouse aorta in the absence or presence of the adventitial (Adv) layer. The housekeeping gene *hypoxanthine guanine phosphoribosyltransferase (hprt)* was used as a control. Data are expressed as the Log2 fold change in mRNA levels relative to murine aortic SMCs in culture and are the mean ± SEM of three aortic specimens, #p≤ 0.05 vs MOVAS SMCs. **d.** Flow cytometry analysis of vSCs isolated from AA and TA regions of the mouse aorta and grown in maintenance media for S100β (dark green) and Sca1 (light green) expression. NE-4C and CH3-10T1/2 cells were used as positive controls, respectively. Filled dark and light green curves represent negative control samples. **e,f.** Relative levels of **(e)** *Myh11* and **(f)** *Cnn1* in vSCs isolated from AA and TA regions of the mouse aorta, compared to AA and TA aortic tissue. Data are expressed as the Log2 fold change in mRNA levels relative to neural stem cells (NE-4C) in culture and are the mean ± SEM of three independent cultures and three aortic specimens, #p≤ 0.05 vs NE-4C cells. **g,h.** Representative immunocytochemical analysis of stem cell markers S100β, Nestin, Sox10, Sox17, Sca1 and SMC differentiation markers, *Cnn1* and *Myh11* in vSCs isolated and grown in maintenance media (MM) from **(g)** AA and **(h)** TA regions of the mouse aorta. Scale bar 20μm. **i.** Absolute telomere length of vSCs compared to cultured MOVAS SMC and fresh aortic tissue. The telomere primer set recognizes and amplifies telomere sequences. The single copy reference (SCR) primer set recognizes and amplifies a 100 bp-long region on mouse chromosome 10, and serves as reference for data normalization. Data are expressed as absolute telomere length (kb) and are the mean ± SEM of three plates and aortic specimens, #p≤ 0.05 vs MOVAs SMCs. **j,k.** Relative levels of Gli1, *Myh11* and *Cnn1* within vSCs isolated from **(j)** AA and **(k)** TA regions of the mouse aorta in the absence or presence of rSHh (0.5 μg/ml) with or without the smoothened inhibitor, cyclopamine (10μM). Data are expressed as the Log2 fold change in mRNA levels relative to vSCs alone (control) and are the mean ± SEM of three representative wells from two independent experiments, #p≤0.05 versus rSHh alone. **l-m.** ChiP analysis of the ratio of (l) H3K4me2:H3K27me3 and (m) H3K27me3:H3K4me2 enrichment at the *Myh11* locus in AA vSCs in the absence or presence of rSHh (0.5 μg/ml) with or without cyclopamine (10μM) for 7 d. **n.** Fold enrichment of H3K4me2 at the *Myh11* locus in AA vSCs in the absence or presence of rSHh (0.5 μg/ml) with or without cyclopamine (15μM) for 7 d. Fresh aortic tissue and mouse ECSs was used as positive and negative controls, respectively Data are the mean ± SEM, n=3, #p≤ 0.05 versus aorta/ESC, *p≤ 0.05 vs vSC, §p≤ 0.05 vs rSHh. **o.** Representative immunocytofluorescence staining and **p**. the fraction of *Cnn1* and *Myh11* positive cells in the absence or presence of rSHh (0.5 μg/ml) with or without cyclopamine (15μM) for 7 days. Representative images shown, Scale bar = 50 μm. Cumulative data is the mean ± SEM, n=3, #p≤ 0.05 versus control, *p≤ 0.05 vs rSHh.

Resident vascular stem cells (vSCs) were subsequently isolated from mouse aortic arch (AA) and thoracic aorta (TA) by enzymatic digestion and sequential plating, and were characterised by flow cytometry, immunocytochemistry and gene expression for both stem cell markers and SMC differentiation markers. Flow cytometric analysis confirmed that the cells were Sca1 and S100β positive when compared to Sca1 and S100β positive control cell lines (i.e., C3H 10T1/2 and NE-4C, respectively) [Figure 5d]. The vSCs did not enrich for *Myh11* or *Cnn1* transcripts when compared to fresh aortic tissue [Figure 5e, f] or express these proteins [Figure 5g,h] but exhibited greater teleomere length as a measure of stemness when compared to freshly isolated aortic SMC and to the immortalised MOVAS SMCs in culture [Figure 5i]. Immunocytochemical analysis revealed that vSCs from both AA and TA regions were Sca1 positive and expressed neuroectodermal markers S100β, Sox10, Sox17, and Nestin but were negative for SMC differentiation markers *Cnn1* and *Myh11* [Figure 5g, h]. The vSCs were enriched for neuroectodermal marker genes *S100β, Sox10 and Nestin* [Suppl Figure 5a-c], but not for mesoderm markers *Kdr, Pax1 and Tbx6* [Suppl Figure 5d-f], when compared to SMCs and NE-4 cells in culture. Collectively, these data indicate that neuroectodermal S100β/Sca1 cells are primarily located within the adventitial perivascular region of atheroprone regions of the mouse aorta and can be isolated and expanded *In vitro* to retain these phenotypic markers.

### Hedgehog signalling promotes myogenic differentiation of resident murine S100β/Sca1 vascular stem cells *In vitro*

The functional effect of sonic hedgehog (SHh) signalling on the growth and myogenic differentiation of vSCs was evaluated *In vitro* using multipotent Sca1/S100β cells isolated from mouse aorta. Whole vessel Gli1 mRNA levels increased over time following ligation [Suppl Figure 6c]. Treatment of vSCs isolated from AA and TA with recombinant SHh (rSHh, 0.5 μg/ml, 24h) promoted Hh signalling by increasing Hh target gene mRNA levels (*Gli 1*, *Gli 2* and *Ptch1*), an effect blocked following pre-treatment with cyclopamine [Figure 5j, k and Suppl Figure 6a, b]. Concurrent treatment of cells with rSHh for 7 days resulted in a significant increase in SMC differentiation marker gene expression (*Cnn1* and *Myh11*), an effect attenuated by pre-treatment of cells with cyclopamine [Figure 5j, k]. SHh treatment also significantly increased the growth of AA vSCs after 7 days, an effect attenuated by pre-treatment with cyclopamine [Suppl Figure 6d].

The level of enrichment of the SMC specific epigenetic histone mark, di-methylation of lysine 4 on histone H3 (H3K4me2) and the repressor mark, tri-methylation of lysine 27 on histone H3 (H3K27me3) at the **Myh11** locus was assessed using chromatin immunoprecipitation (ChIP) assays before and after rSHh treatment. Freshly isolated differentiated aortic medial SMC and murine embryonic stem cells (mESCs) were used as positive/negative controls for each mark and were markedly enriched for H3K4me2 and H3K27me3 at the **Myh11** promoter, respectively [Figure 5l-n]. In contrast to medial SMCs, AA vSCs were enriched for H3K27me3 but lacked enrichment of H3K4me2 at this locus. Treatment with recombinant SHh (rSHh, 0.5 μg/ml, 7d) increased the enrichment levels of the H3K4me2 mark while concurrently decreasing the enrichment of the H3K27me3 mark, an effect attenuated by cyclopamine [Figure 5l-n]. Finally, SHh treatment significantly increased the fraction of *Cnn1*^+^ and *Myh11*^+^ cells in AA vSCs, an effect attenuated by pre-treatment with cyclopamine [Figure 5o,p].

### S100β cells populate human arteriosclerotic lesions

The expression of SMA, S100β and the hedgehog receptor, Ptch1, was determined by immunofluorescence staining in normal and arteriosclerotic human aortic sections. The expression of SMA was abundant in the medial layers of both normal and arteriosclerotic vessels while reduced within the neointima [Figure 6a]. The expression of S100β was confined to the adventitial layer of normal vessels but was significantly enhanced within the adventitial and intimal layers of arteriosclerotic sections [Figure 6a]. The expression of Ptch1 was confined to the adventitial layers of normal vessels but was also sparsely expressed in the medial and intimal layers of arteriosclerotic sections [Figure 6a].

**Figure 6.**
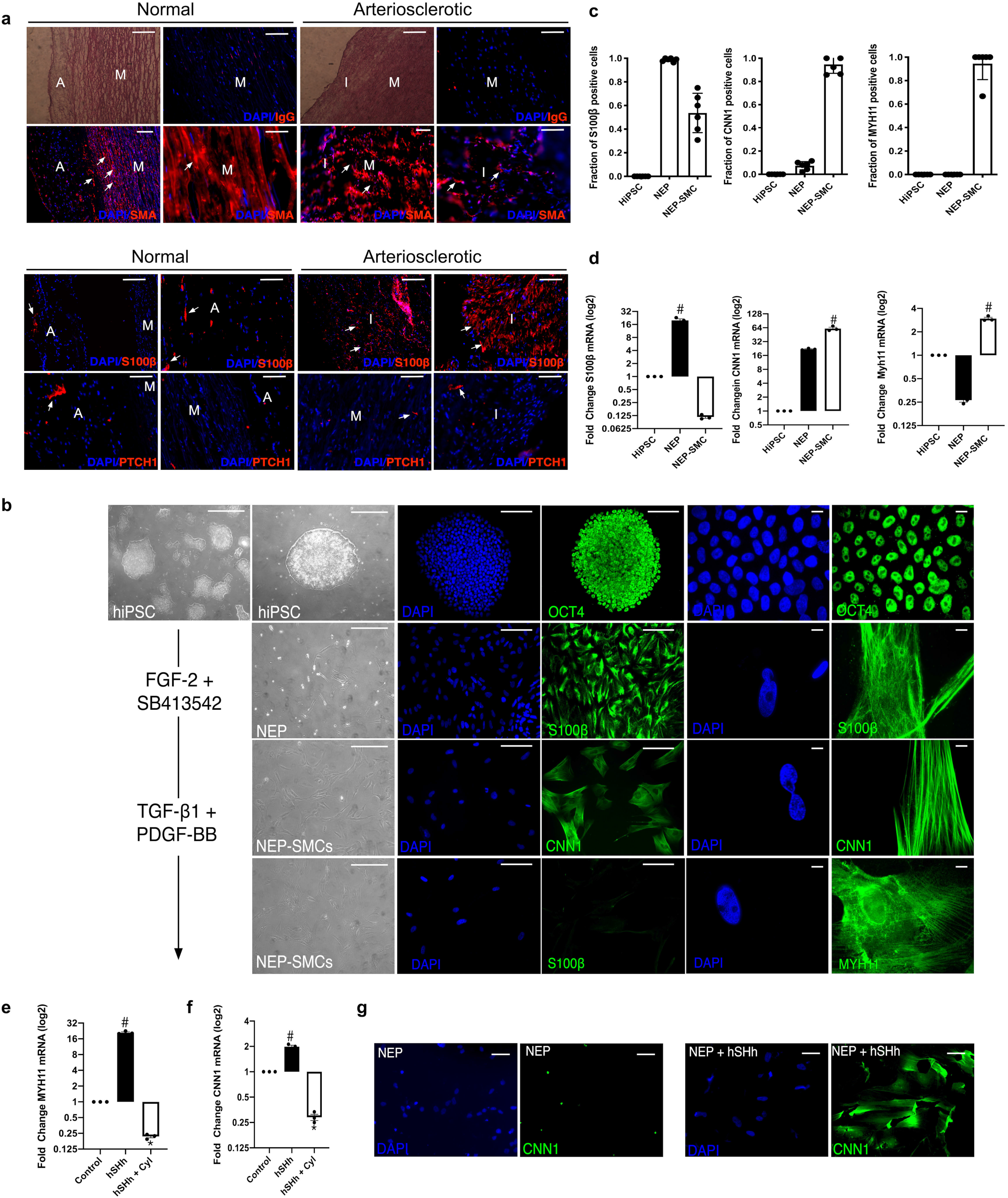
Human neuroectoderm progenitors express S100β cells and are present in vascular lesions. **a.** Representative phase contrast photomicrographs, hematoxylin and eosin (H&E) staining, DAPI staining (blue) and immunocytochemical staining for smooth muscle cell α-actin (SMA) (red), S100β (red) and Ptch1 as indicated in the adventitial (A), medial (M), and intimal (I) sections of normal human and arteriosclerotic arteries. Scale bar=50μm. Data are representative of 3 sections from 3 patient cohorts. **b.** Phase contrast images, fluorescence staining of DAPI nuclei (blue) and immunoctytochemical analysis of the expression of OCT4 (green), S100β (green) in NEPs and Cnn1 (green) and Myh11 (green) in NEP-derived SMCs. All cells were generated as described previously. Scale bar = Right panels: 50μm, Left panels: 20μm. Data are representative of five images per experimental group from three independent cultures. **c.** The fraction of S100β, Cnn1^+^ and Myh11^+^ cells in HiPSCs, NEPs and NEP-SMCs. Data are mean ± SEM, n=3, #p≤ 0.05 versus HiPSCs. **d.** Relative levels of S100β, Cnn1 and Myh11 in HiPSCs, NEPs and NEP-SMCs, respectively. Data are expressed as the Log2 fold change in mRNA levels relative to HiPSCs in culture and are the mean ± SEM, n=3, #p≤ 0.05 versus HiPSCs. **e,f.** Relative levels of (e) Cnn1 and (f) Myh11 in NEPs in the absence or presence of hSHh (0.5μg/ml) with or without cyclopamine (10μM) for 12 d. Data are representative of three independent cultures, #p≤ 0.05 versus NEPs, *p≤ 0.05 vs hSHh. **g.** Immunoctytochemical analysis of Cnn1 (green) expression and DAPI (blue) staining in NEPs in the absence or presence of recombinant hSHh (0.5 μg/ml) for 12 d. Data are representative of five images per experimental group from two independent cultures.

### Hedgehog signalling promotes myogenic differentiation of human induced pluripotent stem cell (HiPSC)-derived neuroectodermal S100β vascular stem cells *In vitro*

To address whether human neuroectoderm S100β progenitors (NEPs) respond to hSHh by undergoing myogenic differentiation, S100β NEPs were isolated from human induced pluripotent stem cells (HiPSC) using a chemically defined protocol ^43^. Human iPSCs enriched for Oct4 were treated with neuroectodermal inductive stimuli (FGF-2 and the selective inhibitor of ALK5, SB413642) for 7 days [Figure 6b] and expressed S100β but not SMC differentiation markers Cnn1 and Myh11 [Figure 6b]. Treatment of NEPs with myogenic media supplemented with TGF-β1/PDGF-BB significantly increased the fraction of Cnn1 and Myh11 cells after a further 12 d in culture, while concomitantly decreasing the number of S100β cells [Figure 6c]. The levels of S100β, Cnn1 and Myh11 transcripts mirrored the changes in protein, where NEP-derived SMCs became significantly enriched for Cnn1 and Myh11 following TGF-β1/PDGF-BB treatment as they diminish their expression of S100β [Figure 6d]. Treatment of NEPs with rSHh (1.0 μg/ml) for 12 d resulted in a significant increase in Cnn1 and Myh11 mRNA levels, an effect attenuated by cyclopamine [Figure 6e, f], concomitant with an increase in the number of Cnn1 positive cells [Figure 6g]. SHh treatment also significantly increased the growth of S100β NEPs after 12 days in culture, an effect attenuated by pre-treatment with cyclopamine [Suppl Figure 6e].

## Discussion

In this study, we used transgenic mice and genetic lineage tracing approaches to delineate the involvement of Hh-responsive S100β resident vSCs in progressing neointima formation following flow restriction. Our findings indicate that intimal thickening involves significant accumulation of Hh-responsive S100β-derived progeny. Elucidation of this new cellular S100β source for generating SMC-like cells following injury expands our current understanding of vascular pathology and may represent an important therapeutic strategy for combating subclinical atherosclerosis.

SMC-like cells within neointimal lesions reportedly are monoclonal/oligoclonal in origin arising from the expansion of a pre-existing resident clonal population of cells ^44–47^. Despite extensive research, their origin has remained controversial ^48^. As a general rule, SMCs are purported to switch from a contractile phenotype to a less mature synthetic phenotype following vascular injury resulting in the loss of expression of SMC differentiation contractile markers, increased proliferation, and the synthesis and release of pro-inflammatory cytokines, chemotaxis-associated molecules, and growth factors ^49^. However, whether these clonal progeny within lesions are derived primarily from differentiated medial SMCs ^8,9,49^, endothelial cells ^10,50^, and/or resident vSCs^10–13^ remains unclear. Over the last decade, vascular SMC lineage tracing studies have provided compelling evidence of a significant role for pre-existing medial SMCs in vascular pathologies ^8,51–53^. Single cell RNA sequence analysis (scRNA-seq) further identified a rare Sca1^+^ Myh11-Cre marked SMC subpopulation that gives rise to functionally distinct neointimal cells during lesion progression in mice ^8,9^. However, although the Myh11-CreERT2 transgene was originally deemed specific for vascular SMC ^54^, data indicating expression of this and other ‘SMC differentiation genes’ in non-SMC populations has recently emerged ^55–57^ raising the possibility that other resident cells may be inadvertently marked and contribute to lesion formation. Myh11-CreER SMC lineage tracing studies have also demonstrated that the fraction of SMCs originating from pre-existing SMCs may be considerably smaller after severe transmural injury in mice further suggesting a contributory role for cells other than medial SMCs ^58^.

In this study, we provide genetic evidence that vascular lesions contain an abundance of dual labelled Hh responsive S100β/Sca1 vSCs derived from a non-SMC perivascular and/or medial S100β parent population. Notwithstanding the fact that Sca1 cells are present in bone-marrow ^59^, bone-marrow cells were not marked using the S100β-CreERT2-tdT. It is therefore most likely that Sca1/S100β cells marked with S100β-CreERT2-tdT within lesions were locally derived ^47^. S100β cells predominate within the adventitial layer of healthy vessels (murine and human) since removal of the adventitia dramatically reduced aortic S100β transcript levels in mice. Nevertheless, a small fraction of S100β/Sca1^+^ cells are present in medial layer of normal vessels and could be successfully isolated from aortic medial segments following enzymatic digestion and sequential plating *In vitro*. Indeed, previous studies using in situ immunohistochemical analysis and scRNAseq also confirms the presence of a sparse Sox10/S100β cell population within the media, in addition to the adventitia ^9,60,61^. However, these enrichment levels were minimal compared to an S100β^+^ neuronal cell and vascular adventitial ‘fibroblast-like cell’ phenotypes within the vessel wall, supporting the contention that a perivascular adventitial S100β cell may be the predominant source of S100β within the vessel wall. Similarly, a discrete population of immature medial cells expressing CD146 that emerge during embryonic development may also contribute to neointimal formation in response to injury ^58^. Whether these CD146 immature SMC progenitors are also S100β/Sca1 remains unclear although S100β vSCs derived from rat and murine aortic explants express CD146 ^12,42^. In the future, it will be important to determine whether medial S100β vSCs differ from CD146 ^58^ and Myh11-CreER Sca1 cells ^9^ in their propensity to modulate different phenotypes and undergo clonal expansion using various injury models.

Adventitial progenitors have long been implicated in the progression of neointimal lesions following various vascular injuries ^66,67^. However, in the absence of Sca1 or c-kit^+^ stem cell lineage tracing experiments *in vivo*, the hypothesis of a regenerative function for adventitial Sca1 or c-kit^+^ vSCs in lesion formation has been difficult to prove until recently. While perivascular cells must first traverse the internal and external elastic lamina to accumulate within the intima following injury ^64^, disruption of the elastic lamellae and flow-dependent enlargement of fenestrae can facilitate such migration ^65–67^. Cell fate mapping studies have now clarified the role for adventitial PDGFR⍰ Sca1 progenitors in lesion progression following severe injury ^11^ while c-kit progenitors minimally facilitate endothelial regeneration or SMC differentiation to SMCs ^68^. As Sca1-CreER marked cells were not present in vascular lesions following wire-induced injury ^11^, the origin of Sca1 neointimal cells following flow restriction in the current study is unlikely to be from a Sca1 parent population despite expressing Sca1 (albeit using a different injury model). Indeed, the percentage of perivascular S100β-CreER marked cells that were Sca1^+^ in normal vessels was < 50% indicating that a Sca1^neg^/S100β^pos^ population might be responsible. In this context, S100β has been shown to maintain an intermediate state of Sca1 progenitor cells following lesion formation^20^. Alternatively, this residual Sca1^pos^/S100β^neg^ population of adventitial Sca1 cells might represent Sca1 cells derived from differentiated Myh11-CreER SMCs that expand in number following injury and contribute to adventitial remodelling^14^ and lesion formation ^9^. While Sca1/S100β cells were predominantly found in the developing intimal layer after 7, 14 and 21 days, these cells were also present in the medial layer following injury. In agreement with previous studies demonstrating significant apoptosis of medial SMCs following flow restriction^69^, the appearance of medial Sca1/S100β cells marked with S100β-CreERT2-tdT following early expression of the pro-apoptotic protein Bax within the media is consistent with regeneration of medial SMCs following injury.

Previous lineage tracing studies have also implicated endothelial cells (EC) that undergo endothelial-to-mesenchymal transition (EndMT) to SMC-like cells as contributing to neointimal formation during vein graft remodelling ^10,50^. EndMT promotes neointimal hyperplasia and induces atherogenic differentiation of EC ^70^. In the current study, EC were not marked with S100β-CreERT2-tdT prior to flow restriction suggesting that the S100β-CreERT2-tdT marked cells within lesions do not originate from an EC parent population. These apparent differences may result from the use of different models of vascular injury where vein graft and atherosclerosis models involve different biomechanical stimuli and/or a different inflammatory environments compared to the flow restriction model ^50,71^. While EC did not contribute to neointimal formation in developed lesions, a potential role for adluminal Sca1 cells in regenerating the endothelium following ligation was observed. As flow restriction leading to alterations in shear stress and cyclic strain is strongly associated with increased endothelial cell apoptosis *In vitro* ^72^ and *in vivo* ^73^, significant Bax expression at the adluminal surface was observed after flow restriction. Bcl-2/Bax ratios are an important rheostat to determine the incidence of apoptosis in general and increases in Bax expression have been widely reported and validated using the TUNNEL assay to indicate apoptosis of endothelial cells *In vitro* in response to low shear stress (LSS) ^74^ and *in vivo* following carotid artery ligation in wildtype mice ^75^ or following flow perturbation in ApoE deficient mice ^76^. The increase in Bax expression concomitant with an increase in the number of Sca1 cells that maintain a CD31 and eNOS endothelial cell phenotype at the adluminal surface is consistent with the appearance of new endothelial cells in the injured vessel that are derived from the proliferation of adluminal Sca1 cells. Indeed, recent cell fate mapping studies using Sca1-Cre-ERT2-tdT mice have since confirmed this observation by demonstrating expansion of Sca1-CreERT marked cells in the intimal layer that maintain CDH5 endothelial cell fate following injury ^11^.

There are many potential reasons for the different outcomes of lineage tracing studies that addressed the origin of lesional SMC-like cells including technical considerations around fixation, tissue preservation, brightness and contrast settings for image acquisition, autofluorescent material such as lipofuscin and single cell resolution ^77^. Another important reason may involve the use of tamoxifen (Tm) in time-sensitive Tm-dependent Cre-LoxP models. Tm-induced Cre recombinase has been shown to indelibly mark cells up to 4 weeks after the last Tm injection thereby potentially confounding the interpretation of many previous studies ^41^. In this context, we found that the fraction of S100β-CreER (tdT) marked cells within the various compartments of ligated vessels four weeks after the last Tm injection was similar to that observed after one week, indicating that, at least in our hands, medial SMC that might undergo de-differentiation and acquire the S100β promoter are not being inadvertently marked after injury.

Several GWAS studies have implicated HHIPL1 at the chromosome 14q32 CAD locus ^27,78^ and identified it as a secreted Hh signaling protein that modulates atherosclerosis-relevant vascular phenotypes ^28^. Here we provide direct evidence that inhibition of Hh signalling controls neointimal thickening following flow restriction by attenuating the volume and accumulation of Sca1 cells within lesions. Furthermore, Hh signalling directly promotes Sca1/S100β cells from mouse and S100β neuroectoderm stem cells derived from human induced pluripotent stem cells (HiPSC) to undergo growth and myogenic differentiation to SMC-like cells *In vitro*. The possibility that Hh responsive S100β/Sca1 cells isolated from murine AA and TA regions are derived from a *Myh11* medial SMC population was ruled out by examining the H3K4me2 epigenetic profile of S100β/Sca1 cells at the *Myh11* locus, which is preserved during SMC phenotypic modulation *In vitro* and *in vivo* following injury^79^. Hh responsive S100β/Sca1 vSCs do not have this SMC histone mark and only become enriched following myogenic differentiation to SMC-like cells, an effect attenuated by the Hh inhibitor, cylopamine. The accumulation of Sca1/S100β cells within lesions may be due, in part, to asymmetric division of Sca1/S100β progenitor stem cells (from the adventitia and/or media) and subsequent proliferation and migration of their partially differentiated myogenic progeny to the intima. Indeed, SHh promotes Sca1/S100β growth and the proliferation of modulated, partially differentiated SMCs *In vitro* ^32^. Moreover, previous studies have shown that it is a Sca1 adventitial population in particular that proliferate following injury using immunohistochemical co-staining with Ki-67 ^31^. This is not surprising as adventitial Sca1 stem cells co-localise with SHh and its receptor, *Ptch1* ^35^ while several previous studies support a role for Hh signalling in vascular lesion formation in mice ^29,38,39^. It appears the Yes-Associated Protein (YAP) is required for incorporation and expansion of Sca1-derived neointimal cells following injury ^11^ where the Hh pathway reportedly acts upstream of the Hippo pathway to regulate stem cell maintenance ^80^. Indeed, our data suggest S100β cells are also present in normal vessels within the adventitial layer where Hh components reside ^81,82^. In arteriosclerotic lesions, S100β expression is exaggerated within the neointimal layer with sparse staining for *Ptch1* present. As S100β NEP-derived SMCs are considered crucial to the generation of SMC in vascular lesions within atheroprone regions of the murine vasculature ^4,43^, it is tempting to speculate that Hh responsive S100β-derived SMC-like progeny may also play a pivotal role in lesion formation in humans. Future studies that address the interactions between Hippo pathway and Hh may yield therapeutic strategies aimed at interfering with YAP activity through inhibition of hedgehog pathway, which downregulates the YAP protein^83,84^.

In conclusion, the identification of a population of S100β/Sca1 cells within vascular lesions that are derived from a non-SMC S100β parent population is novel and highlights these cells as an important source for intimal thickening following injury. Furthermore, the role of SHh in mediating this process expands our understanding of the mechanisms contributing to the progression of vascular lesions. While Hh signalling has previously been implicated in the development of vascular lesions through modulation of a differentiated SMC phenotype ^28^, our study further suggests that Hh signalling may also control myogenic differentiation and accumulation S100β/Sca1 derived SMC-like progeny within vascular lesions. While hedgehog signalling appears critical, further research is required to investigate the precise mechanisms driving migration, proliferation and differentiation of stem cells during lesion formation. It is possible that SMC-like cells within lesions comprise several different stages of the myogenic, osteogenic and myeloid differentiation of various stem cells and hence display a variety of divergent phenotypes. The elucidation of these mechanisms will provide vital information needed for the development of more efficient and novel therapies for vascular proliferative diseases. Our results provide a potential strategy for the treatment of atherosclerosis by combating cell migration and asymmetric differentiation of a perivascular-derived S100β parent vSC population.

## Methods

### Mice Breeding and Genotyping

All procedures were approved by the University of Rochester Animal Care Committee in accordance with the guidelines of the National Institutes of Health for the Care and Use of Laboratory Animals. Sca1-eGFP transgenic mice were obtained from JAX labs; Stock #012643, strain name B6.Cg.Tg(Ly6a-EGFP)G5Dzk/j. These transgenic mice have an enhanced green fluorescent protein (eGFP) under the control of murine lymphocyte antigen 6 complex, locus A (Ly6a) promoter. Hemizygous Ly6a-GFP mice are viable, fertile, normal in size and do not display any gross physical or behavioural abnormalities ^85^.

S100β-EGFP/Cre/ERT2 transgenic mice (JAX Labs, stock #014160, strain name B6;DBA-Tg(S100β-EGFP/cre/ERT2)22Amc/j) express the eGFPCreER^T2^ (Enhanced Green Fluorescent Protein and tamoxifen inducible cre recombinase/ESR1) fusion gene under the direction of the mouse S100β promoter. Ai9 mice (Jax Labs, stock #007909, strain name B6.Cg-Gt(ROSA)26Sor^tm9(CAG-tdTomato)Hze^ /J) express robust tdTomato fluorescence following Cre-mediated LoxP recombination. For lineage tracing experiments S100β-eGFP/Cre/ERT2– dTomato double transgenic mice of both genders were generated by crossing S100β-eGFP/Cre-ERT2 mice with Ai9 reporter mice. The tdTomato transgene expression pattern corresponds to genomic marked S100β cells, and the eGFP transgene expression pattern corresponds to constitutive expression of S100β. Mice were genotyped using genomic DNA prepared from tail samples. All male and female mice were included in the study and were 8-10 weeks old.

### Tamoxifen-induced genetic lineage tracing

S100β-CreER-Rosa26tdT mice (average weight 20g, 6-8 wks old) were injected IP with tamoxifen (Tm) dissolved in corn oil at 75 mg/kg for 5 consecutive days. Carotid artery ligation (partial, or complete), or sham operation, was performed at 1 wk or 4 wks after the last injection of Tm. At the indicated time post-ligation or sham operation, anesthetized mice were perfusion fixed and carotid arteries harvested for analysis.

### Carotid Artery Ligation

Ligation of the left common carotid artery was performed essentially as described previously ^38^ one and four weeks after Tm-induced cre-recombination, in male and female S100β-eGFP/Cre/ERT2–dTomato double transgenic mice. Prior to surgery mice received a single dose of Buprenorphine SR (sustained release) analgesia (0.5-1.0 mg/kg SQ) (and every 72 hrs thereafter as needed). The animal was clipped and the surgical site prepped using betadine solution and alcohol. A midline cervical incision was made. For partial carotid artery ligation, with the aid of a dissecting microscope, the left external and internal carotid arterial branches were isolated and ligated with 6-0 silk suture reducing left carotid blood flow to flow via the patent occipital artery. The neck incision (2 layers, muscle and skin) was sutured closed. Partial ligation of the left carotid artery in this manner resulted in a decrease (∼80%) in blood flow, leaving an intact endothelial monolayer. Buprenorphine was administered at least once post-op 6-12 hrs. For complete ligation, the left common carotid was isolated and ligated just beneath the bifurcation with 6-0 silk suture. In sham surgery group, carotid was similarly manipulated but not ligated. The neck incision (2 layers, muscle and skin) was sutured closed and the animal allowed recover under observation. After 3, 7, 14 and 21 days, mice were perfused with 4% PFA via the left ventricle for 10 min, and carotid arteries were harvested for analysis.

### Cyclopamine treatment

Sca1-eGFP mice were treated with the smoothened inhibitor, cyclopamine, or the vehicle 2-hydropropyl-β-cyclodextrin (HβCD) (Sigma-Aldrich) alone as a control, (essentially as described previously) ^86^. Mice were injected (250 μl max volume) intraperitoneally (IP) 1 day before ligation, then every other day after ligation at a dose of 10 mg/kg Cyclopamine (Sigma-Aldrich) dissolved in a solution of 45% (w/v) HβCD. There was no effect of cyclopamine treatment on mouse body weight, compared to the HβCD vehicle control group (data not shown).

### Histomorphometry

At specified times post-ligation, mice were anesthetized (ketamine/xylazine) and perfusion fixed with 4% paraformaldehyde in sodium phosphate buffer (pH 7.0). Fixed carotids were embedded in paraffin for sectioning. Starting at the carotid bifurcation landmark (single lumen) a series of cross-sections (10 × 5 μm) were made, every 200 μm through 2 mm length of carotid artery. Cross-sections were de-paraffinized, rehydrated in graded alcohols and stained with Verhoeff-Van Gieson stain for elastic laminae and imaged using a Nikon TE300 microscope equipped with a Spot RT digital camera (Diagnostic Instruments). Digitized images were analyzed using SPOT Advanced imaging software. Assuming a circular structure *in vivo*, the circumference of the lumen was used to calculate the lumen area, the intimal area was defined by the luminal surface and internal elastic lamina (IEL), the medial area was defined by the IEL and external elastic lamina (EEL), and the adventitial area was the area between the EEL and the outer edge, essentially as described by us previously ^38^

### Haematoxylin and Eosin Staining of Tissue

For human artery cross sections, slides were placed in ice-cold acetone and held at −20°C for 15 min. They were then washed 3 × PBS before immersion in Harris Haematoxylin at room temperature for 8 min. Slides were processed by sequential washing – in distilled water for 5 min; 1 % acid alcohol for 30 sec; 0.1 % sodium bicarbonate for 1 min and 95 % ethanol. Slides were then washed in Eosin for 1 min, 75 % ethanol for 3 min, 95 % ethanol for 3 min and 100 % ethanol for 3 min before being submerged in Histoclear for 3 min and permanently mounted using 3 drops of DPX, cleaned with 100 % ethanol before analysis.

For murine vessels, paraffin rehydration was conducted at room temperature by immersing the slides in xylene for 20 minutes. The slides were then immersed in the following gradients of ethanol for 5 minutes respectively: 100 %, 90 %, 70 % and 50 % ethanol. The slides were then rinsed in distilled H_2_O_2_ before washing in 1x PBS for 10 minutes. The slides were stored in 1x PBS until ready when they were immersed in Harris Haematoxylin at room temperature for 8 min.

S100β^+^ cells were identified as either “genomically marked” red tdTomato expressing cells or “currently expressing” green fluorescent protein (eGFP) cells visualized in deparaffinized S100β-eGFP/Cre/ERT2–dTomato mouse carotid cross sections mounted with Sigma Fluoroshield with DAPI, using an FV1000 Olympus or a Nikon A1R HD laser scanning confocal microscope. Numbers of red or green fluorescent cells and DAPI nuclei (blue) in carotid cross section images from different experimental groups were either analyzed by Fiji ImageJ software in each vessel compartment using a grid system.

### Whole mount histochemical analysis

Briefly, arteries harvested from transgenic mice were washed in PBS before transfer to 10 % neutral buffered formalin and incubated at 4 °C overnight. The tissue was rinsed 3 × PBS before mounting on histology slides using DAPI containing fluoroshield (Sigma). The whole tissue was then imaged using a Nikon Eclipse TS100 inverted microscope, EXFO X-Cite™ 120 fluorescence system and assessed using SPOT imaging software.

### Immunofluorescent Staining of tissues

Immunostaining essentially as previously described ^38^. Carotid artery cryosections were air-dried for 1 h at room temperature, followed by incubation with blocking buffer (5% donkey serum, 0.1% Triton X-100 in PBS) for 30 minutes at room temperature and then incubated with primary antibody overnight at 4°C in antibody solution (2.5% BSA, 0.3M Glycine and 1% Tween in DPBS). Murine and human arterial sections were stained with primary antibodies [Supplementary Table I]. Isotype IgG control and secondary antibody only controls were performed. For antigen retrieval, slides were brought to a boil in 10 mM sodium citrate (pH 6.0) then maintained at a sub-boiling temperature for 10 minutes. Slides were cooled on the bench-top for 30 minutes then washed in deionized water (3 × 5 min) each before being washed in PBS (3 × 5 min). The antigen retrieval protocol diminishes endogenous eGFP and tdT tomato transgene signals. Therefore, those sections were co-stained with anti-eGFP antibody and anti-Td tomato antibody [Supplementary Table I].

For immunofluorescence staining, 5 consecutive images were obtained and processed using ImageJ software™ to analyze the collected images. Mouse carotid artery cross sections were mounted with fluoroshield with DAPI and analyzed using an FV1000 Olympus or a Nikon A1R HD laser scanning confocal microscope. Images were merged using the Image color-merge channels function. Merged signals and split channels were used to delineate the signals at single-cell resolution. Settings were fixed at the beginning of both acquisition and analysis steps and were unchanged. Brightness and contrast were lightly adjusted after merging.

### Immunocytofluorescent staining of cells

Cells seeded onto UV sterilized coverslips were fixed with 3.7 % formaldehyde, (15 min, RT). If cells required permeabilization for the detection of intracellular antigens, cells were incubated in 0.025 % Triton X-100 PBS (room temp, 15 min). All coverslips were blocked (1 hr, RT) using 5 % BSA, 0.3 M Glycine, 1 % Tween PBS solution (1 hr, RT). Cells were incubated with primary antibodies overnight at 4°C [Supplemental Table II], then washed twice with PBS to remove any unbound primary antibody before being incubated (1 hr, RT) with the recommended concentration of fluorochrome-conjugated secondary antibodies diluted in blocking buffer [Supplementary Table II]. Following 2x wash in PBS, cell nuclei were stained using DAPI: PBS (dilution 1:1000) (15 min, RT). For each primary and secondary antibody used, a secondary control and an IgG isotype control was performed to assess nonspecific binding. An Olympus CK30 microscope and FCell™ software was used to capture images. Images were analyzed using ImageJ software as described above. Settings were fixed at the beginning of both acquisition and analysis steps and were unchanged. Brightness and contrast were lightly adjusted after merging.

### Cell culture and harvesting

Murine aortic SMCs (MOVAS (ATCC® CRL-2797™) were cultured in Dulbeco’s Modified Eagles growth medium (DMEM) supplemented with Fetal Bovine Serum (10%), L-glutamine (2 mM, Gibco) and Penicillin-Streptomycin (100U/mL, Gibco). Murine neuroectodermal stem cells (mNE-4Cs, ATCC® CRL-2925™) were grown in Eagle’s Minimum Essential Medium supplemented with Fetal Bovine Serum (10%), L-glutamine (2 mM, Gibco) and Penicillin-Streptomycin (100U/mL, Gibco) using Poly-L-Lysine coated plates (15 μg/ml). Murine embryonic C3H 10T1/2 cells (ATCC® CRL-226™) were grown in Eagle’s Basal medium supplemented with heat-inactivated fetal bovine serum to a final concentration of 10% and 2mM L-glutamine. Murine embryonic stem cells (mESCs) ES-D3 [D3] (ATCC® CRL-1934™) were maintained in culture in Mouse ES Cell Basal Medium (ATCC SCRR-2011) supplemented with 2-mercaptoethanol (0.1 mM) and 15% ES-Cell Qualified FBS (ATCC SCRR-30-2020) and a 1,000 U/mL mouse leukemia inhibitory factor (LIF) (Millipore Cat. No. ESG1107). Murine vSCs from AA and TA regions of the mouse aorta were grown in DMEM media supplemented with chick embryo extract (2%), FBS embryonic qualified (1%, ATCC), B-27 Supplement, serum free, N-2 Supplement (Cell Therapy Systems), recombinant mouse bFGF (0.02 μg/mL), 2-Mercaptoethanol (50nM), retinoic acid (100 nM), Penicillin-Streptomycin (100U/mL, Gibco) (MM).

### Isolation of S100β/Sca1 murine resident vascular stem cells (vSCs)

Using an optimised enzymatic cell dissociation technique protocol vSCs from both atheroprone (aortic arch - neuroectodermal, NE), and athero-resistant (thoracic aorta-paraxial mesodermal, PM) regions of the murine aorta were isolated using sequential seeding on adherent and non—adherent plates. Mouse thoracic aortas (4 at a time) were harvested and placed in cold Hank’s solution for adipose tissue removal and lumen rinsing. The adventitia was enzymatically removed by incubation of the vessels in collagenase solution of MEMα containing GlutaMAX− (2 ml), collagenase type 1A (0.7 mg/ml), soybean trypsin inhibitor (50 mg/ml), and bovine serum albumin (1 mg/ml) for approx. 10-20 min at 37°C. Once the adventitia became lose, it was carefully removed as an intact layer using forceps under a dissecting microscope. The aorta was divided into aortic arch (AA) and thoracic aorta (TA) regions and cut into 1 mm pieces and digested with Elastase type III (Sigma) at 37°C. Dispersed cells were centrifuged and washed twice in warm maintenance medium (MM) (DMEM supplemented with 2% chick embryo extract, 1% FBS, 0.02 μg/ml, bFGF basic protein, B-27 Supplement, N-2 supplement, 1% Penicillin-Streptomycin, 50 nM 2-Mercaptoethanol and 100 nM retinoic acid) before seeding (1st seeding) on a 6-well non-adherent plate in MM. At 48 h, suspension cells were transferred (2nd seeding) to a fresh well of a CELLstart™ pre-coated 6-well plate in MM. Additional MM was added to the remaining cells in the non-adherent surface. Cells were incubated at 37°C, 5% CO_2_ for 1 week with minimal disturbance. The stem cells exhibited a uniform neural-like morphology in low density culture adopting a dendritic-like tree shape and retaining their morphological characteristics at low density throughout repeated passage. Cells were fed with MM every 2-3 days and passaged every 3-4 days or when ~70% confluent.

### Generation of human neuroectoderm progenitors (NEPs) from induced pluripotent stem cells (HiPSCs)

Human induced pluripotent stem cells (HiPSC) were obtained from HipSci (Cambridge, UK) and cultured per the manufacturer’s instructions. Briefly, human dermal fibroblasts were isolated from excess skin of patients undergoing plastic surgery and reprogrammed as previously described ^87^. HiPSCs were grown on plates pre-coated using Vitronectin (10 μg/ml) and maintained in complete Essential eight medium replacing 95% of the medium daily. A ROCK inhibitor (10 μM) was added to optimize colony growth. Cells were sub-cultured with 0.05 mM EDTA for 4-8 min at room temperature until colonies displayed bright halos around the edges and small holes appeared throughout. Cells were frozen down in knock-out serum solution with 10% DMSO. HiPSCs at 70% confluency were routinely split at a ratio of 1:10 and cells were used between passage 20-40.

For neuroectoderm differentiation, cells were grown in chemically defined neuroectodermal medium (CDM) containing IMDM, F12 nutrient mix, chemically defined lipid concentrate, transferrin, monothioglycerol, penicillin-streptomycin and 0.5 g of PVA, and insulin, as previously described ^43^. Neuroectoderm differentiation was initiated in CDM supplemented with FGF2 (12 ng/ml) and SB431542 (10 μM) for 7 days. Neuroectodermal (NE) progenitors were maintained in CDM-PVA supplemented with FGF2 (12 ng/ml) and SB431542 (10 μM) and sub-cultured using TrypLE™ for 3–5 min at 37°C in a 5% CO_2_ incubator. NE progenitors were used between passage 1-10.

To initiate myogenic SMC differentiation, NEPs were subcultured in SMC differentiation medium containing CDM + PDGF-BB (10 ng/ml, PeproTech) + TGF-β1 (2 ng/ml, PeproTech) for 12 days. Neuroectoderm-derived SMCs (NE-SMC) were maintained in CDM supplemented with 10% FBS. After 12 d of PDGF-BB and TGF-β1 treatment, all cells appeared spindle-shaped. Complete medium was then changed to DMEM 10% FBS, 1% Penicillin streptomycin for expansion. Cells were frozen down using 90% (volume/volume) FBS + 10% (vol/vol) DMSO. The NE-SMC were used between passage 3-20.

### Human tissue

Tissue sections from paired normal human aorta and arteriosclerotic vessels were obtained from Amsbio LLC, tissue repository.

### Chromatin Immunoprecipitation

ChIP was performed on cultured cells as previously described with slight modifications ^79^. Cells at 70% confluency were fixed with 1% paraformaldehyde (10 min, RT). Cross-link was stopped by addition of 125 mM glycine for 10 min. The cross-linked chromatin was sonicated to shear chromatin into fragments of 200–600 base pairs. The sheared chromatin was immunoprecipitated with 2 μg tri-methyl-histone H3 (Lys27) and di-methyl-histone H3 (Lys4), while negative control was incubated with mouse IgG and input DNA without antibody using the CHIP—IT Express HT Kit from Active Motif (Cat no: 53018) according to the manufacturer’s instructions [Supplementary Table III]. A chromatin IP DNA Purification Kit (Cat No 58002 Active Motif) was used to purify the samples after CHIP before PCR was performed using *Myh11* primers [5’-CCC TCC CTT TGC TAA ACA CA - 3’ and 5’ - CCA GAT CCT GGG TCC TTA CA – 3]. Sensimix SYBR® no-ROX Bioline Kit (QT650) was used to perform Real Time One-Step PCR according to the manufacturers’ instructions. The antibodies used for ChIP are outlined in Supplemental Table I

### Quantitative PCR

Total RNA was prepared from cultured cells using the ReliaPrep™ RNA Cell Miniprep System kit from Promega according to the manufacturer’s protocol. Two micrograms of RNA was used for reverse transcription with Rotor-Gene SYBR Green RT-PCR (QIAGEN) or The SensiMix™ SYBR® No-ROX (BioLine) protocols for Real TimeOne-Step RT-PCR using the Real Time Rotor-GeneRG-3000™ light cycler from Corbett Research using primers listed in Supplementary Table IV.

### Flow cytometry and analysis

Isolated cells were incubated with I fluorochrome-conjugated antibodies and control IgG’s or (ii) the appropriate primary antibody with fluorochrome-conjugated secondary antibodies at a final concentration of 1μg per sample and incubated at 4°C for 30 minutes [Supplementary Table V]. Fluorescence labeled cells analysis was performed using a BD FACS Aria flow cytometry system (BD Biosciences), and data were analyzed with FlowJo™ software (Tree Star, Ashland, Ore) and De Novo software FCS Express 4 Flow Cytometry (Pasadena, CA)

### Telomere Assay

The average telomere length in cells was measured using a ScienCell’s Absolute Mouse Telomere Length Quantification qPCR Assay Kit (AMTLQ) according to the manufacturer’s instructions. The telomere primer set recognizes and amplifies telomere sequences and is designed to directly measure the average telomere length of murine cells.

### QUANTIFICATION AND STATISTICAL ANALYSIS

All data were determined from multiple individual biological samples and presented as mean values ± standard error of the mean (SEM). All *In vitro* experiments were performed in triplicate and repeated three times unless otherwise stated. All mice were randomly assigned to groups (including both male and female), and the analysis was performed blind by two groups. For statistical comparisons, all data was checked for normal gaussian distribution before parametric and non-parametric tests were performed. An unpaired two-sided Student’s t-test was performed using GraphPad Prism software v9™ for comparing differences between two groups, and ANOVA test for over multiple comparisons. Significance was accepted when p≤ 0.05.

## Data Availability

The data, materials and reagents generated or analysed that support the findings of this study are available from the corresponding authors (PAC and EMR) on reasonable request.

## Acknowledgments

We thank Diana Scott for histological tissue processing, and Drs. Linda Callahan and Paivi Jordan for confocal microscope expertise. This research was funded in part by Science Foundation Ireland grant SFI-11/PI/1128, Health Research Board (HRB) of Ireland grant HRA-POR-2015-1315, the European Union’s INTERREG VA Programme, managed by the Special EU Programmes Body (SEUPB) to PAC, and NIH R21AA020365 and R21AA023213 to EMR, Irish Research Council (IRC) GOIPG/2014/43 (M.DiL) and GOIPG/2016/1529 (E.C), King Abdulla Scholarship IR15208 ID1066336148 to Y.G.

## Author Contributions

E.F., W.L., J-C.H, performed the animal experiments and M.DiL and SH performed the blind analysis of the images. E.F performed the human arteriosclerotic tissue analysis. E.F., M.DiL., R.H., D.B., E.C., performed the murine and rat cell culture experiments, D.B performed the HiPSCs cell work. EC performed the telomere assay and R.H.J, MdiL and Y.G., performed the ChIP analysis. E.M.R., C.L., and P.A.C drafted the manuscript, revised the manuscript and discussed the results. E.M.R., and P.A.C., designed and coordinated the experiments, interpreted the data, revised and confirmed the paper. All authors reviewed and confirmed the manuscript.

