## Supplemental Tables and Figures for "The calcium binding protein S100β marks hedgehog-responsive resident vascular stem cells within vascular lesions"

**Supplementary Table I**

Antibodies and their corresponding dilutions used in immunohistochemistry.

| Antibody/Product Name | Supplier/Product Number | Dilutions |
|---|---|---|
| Rat anti-mouse Sca1/Ly6A/E antibody [D7] (FITC) antibody | Abcam (ab25031) | 1/200 |
| Rabbit anti-mouse/rat alpha smooth muscle cell α-actin antibody | Abcam (ab5694) | 1/200 |
| Anti-Patched / PTCH1 antibody, Mouse monoclonal | Abcam (Ab55629) | 1/100 |
| Anti-Actin, α-Smooth Muscle antibody, Mouse monoclonal | Sigma (A5228) | 1/200 |
| Anti-S100-β (CT) Antibody, clone EP1576Y, rabbit monoclonal | Millipore (04-1054) | 1/100 |
| Anti-Gli2 antibody, rabbit polyclonal | Novus Biologicals (NBP2-23602SS) | 1/50 |
| Chicken anti-GFP antibody | Abcam (ab13970) | 1/1000 |
| Rabbit Anti-RFP/dT antibody | Abcam (ab62341) | 1/1000 |

**Supplementary Table II**

Antibodies and their corresponding dilutions used in immunocytochemistry/western blot.

| Antibody/Product Name | Supplier/Product Number | Dilutions |
|---|---|---|
| Mouse anti-mouse/rat nestin [Rat-401] | Abcam (ab11306) | 1/200 |
| Rabbit anti-mouse Calponin [EP798Y] | Abcam (ab46794) | 1/200 |
| Goat anti-mouse/rat/human smooth muscle Myosin heavy chain | Santa Cruz (sc-79079) | 1/200 |
| Mouse anti-human/rat SOX10 | R&D System (MAB2864) | 1/100 |

| | | |
|---|---|---|
| Mouse anti-human SOX17 | R&D System (MAB1924) | 1/100 |
| Rabbit Anti-S100β | Merck Millipore (ABN59) | 1/100 |
| Rabbit Anti-mouse/rat S100β [EP1576Y] | Abcam (ab52642) | 1/100 |
| Rabbit anti-mouse/rat Sca1 | Millipore (AB4336) | 1/100 |
| Alexa Fluor® 488 Goat anti-mouse IgG | Invitrogen (A-11001) | 1/1000 |
| Alexa Fluor® 488 Goat anti-rabbit IgG | Invitrogen (A-11008) | 1/1000 |
| Alexa Fluor® 488 Donkey anti-goat IgG | Invitrogen (A-11055) | 1/1000 |

**Supplementary Table III**

Antibodies used in Chromatin Immunoprecipitation (ChIP)

| Antibody/Product Name | Supplier/Product Number |
|---|---|
| Rabbit anti-mouse Tri-Methyl-Histone H3 (Lys27) [C36B11] | Cell Signalling Technology (9733S) |
| Rabbit anti-mouse Di-Methyl-Histone H3 (Lys4) [C64G9] | Cell Signalling Technology (9725S) |
| Normal Rabbit IgG (ChIP graded) | Cell Signalling Technology (2729) |

**Supplementary Table IV**

Customised primers used in this study from Integrity DNA Technology (IDT).

| Customised primer | | Sequences |
|---|---|---|
| Mm_Sm-mhc (Myh11) | Forward | 5' - GCA GTG AGC TCT CAG TCA TC - 3’ |
| | Reverse | 5' - CAA TGC CTC CTC TGA CAA GT - 3' |
| Mm_Cnn1 | Forward | 5’ - GCT TGT CTG CTG AAG TAA AGA AC - 3' |
| | Reverse | 5’ - TCC ATG AAG TTG TTC CCG ATG - 3' |
| Mm_Gli1 | Forward | 5' - TTG GAT TGA ACA TGG CGT CT - 3' |
| | Reverse | 5' - CCT TTC TTG AGG TTG GGA TGA - 3' |

| | | |
|---|---|---|
| Mm_Hprt | Forward | 5’ - GGC TAT AAG TTC TTT GCT GAC CTG C - 3' |
| | Reverse | 5’ - GCT TGC AAC CTT AAC CAT TTT GGG - 3' |
| Mm_Gapdh | Forward | 5’ - GCC TCC AAG GAG TAA GAA AC - 3' |
| | Reverse | 5’ - GCC TCC AAG GAG TAA GAA AC - 3' |
| Mm_ Sm-mhc (Myh11) for ChIP PCR | Forward | 5’ - CCC TCC CTT TGC TAA ACA CA - 3' |
| | Reverse | 5’ - CCA GAT CCT GGG TCC TTA CA - 3' |

Primers used in this study from QIAGEN.

| Primer | Product Name | Product Code |
|---|---|---|
| mHprt-1 | Mm_Hprt_1_SG QuantiTect Primer Assay | QT00166768 |
| mS100β | Mm_S100β_1_SG QuantiTect Primer Assay | QT00151536 |
| mSox10 | Mm_Sox10_1_SG QuantiTect Primer Assay | QT00295204 |
| mNestin | Mm_Nes_1_SG QuantiTect Primer Assay | QT00316799 |
| mGapdh | Mm_Gapdh_3_SG QuantiTect Primer Assay | QT01658692 |
| mPax6 | Mm_Pax6_1_SG QuantiTect Primer Assay | QT01052786 |
| mPax1 | Mm_Pax1_1_SG QuantiTect Primer Assay | QT01052779 |
| mKdr | Mm_Kdr_1_SG QuantiTect Primer Assay | QT00097020 |
| mTbx6 | Mm_Tbx6_1_SG QuantiTect Primer Assay | QT00098861 |
| hCNN1 | Hs_CNN1_1_SG_QuantiTech Primer | QT00067718 |
| hHPRT1 | Hs_HPRT1_1_SG QuantiTect Primer | QT00059066 |
| hMYH11 | Hs_MYH11_1_SG QuantiTech Primer | QT00069391 |
| hS100β | Hs_S100B_1_SG QuantiTect Primer Assay | QT00059164 |

**Supplementary Table V**

Antibodies and their corresponding dilutions used in Flow Cytometry.

| Antibody/Product Name | Supplier/Product Number | Dilutions |
|---|---|---|
| Rat anti-mouse Sca1 (Ly-6A/E) [E13-161.7] | STEMCELL Technology (60032) | 1/100 |

| | | |
|---|---|---|
| Rat anti-mouse IgG2a, kappa Isotype [RTK2758] | STEMCELL Technology (60076) | 1/100 |
| Rabbit Anti-mouse/rat S100β [EP1576Y] | Abcam (ab52642) | 1/100 |
| Normal Rabbit IgG (F graded) | Cell Signalling Technology (2729) | 1/100 |
| Alexa Fluor 647 Goat anti-mouse (H+L) | Life Technologies (A- A-21235) | 1/100 |

### Supplementary Figures

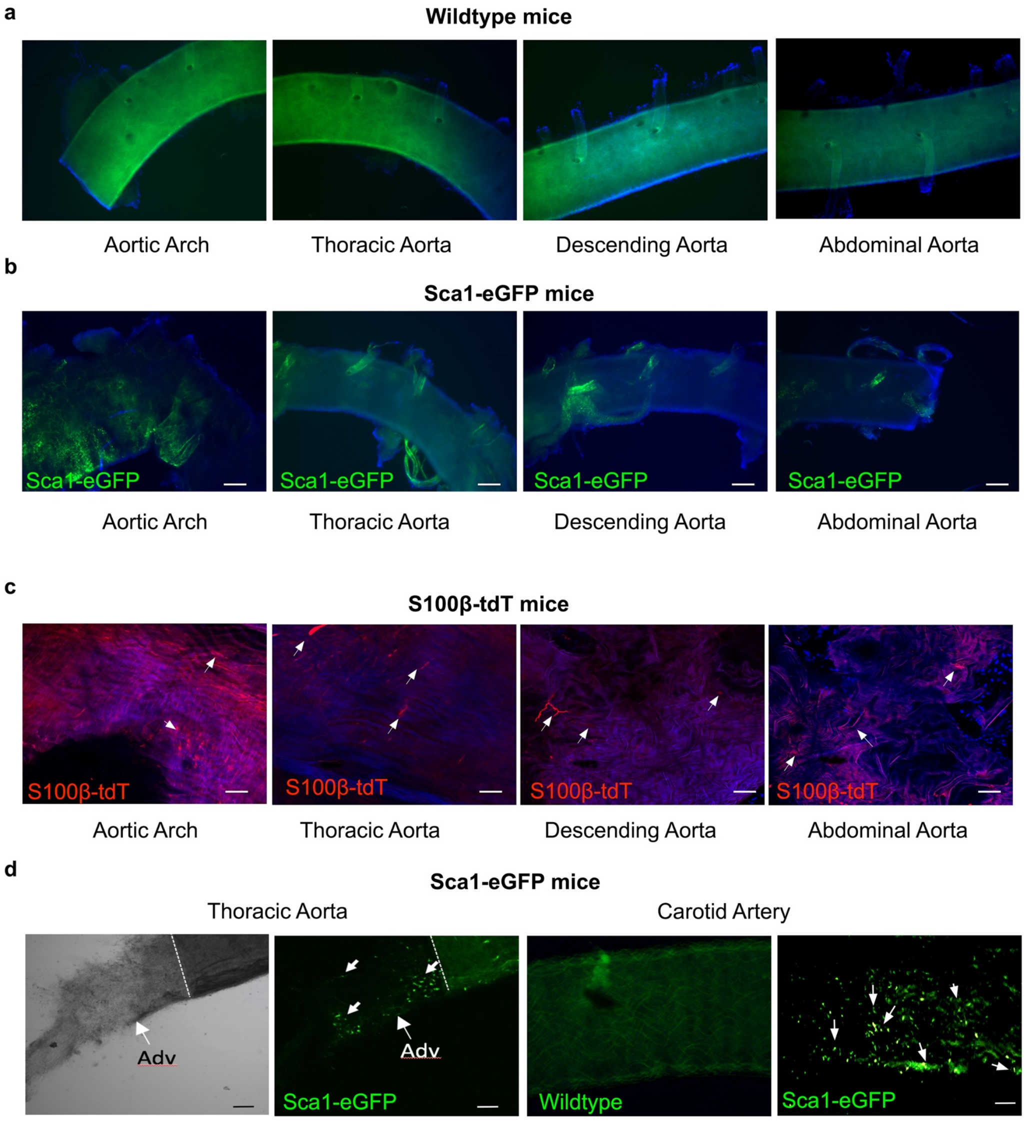


**Suppl Figure 1 (related to Figure 1). Sca1 and S100β cells are present in perivascular regions of murine arteries.**

Whole mount analysis of **a.** wildtype control**, b.** Sca1-eGFP expression and **c.** S100β-Cre-ER2-tdT expression in aortic arch, thoracic aorta, descending aorta and abdominal aorta. **d.** Sca1-eGFP expression in mouse aorta following removal of the aortic media (dashed line) and in carotid artery of Sca1-eGFP transgenic mice. Each image is representative of 5 images from 3 animals.

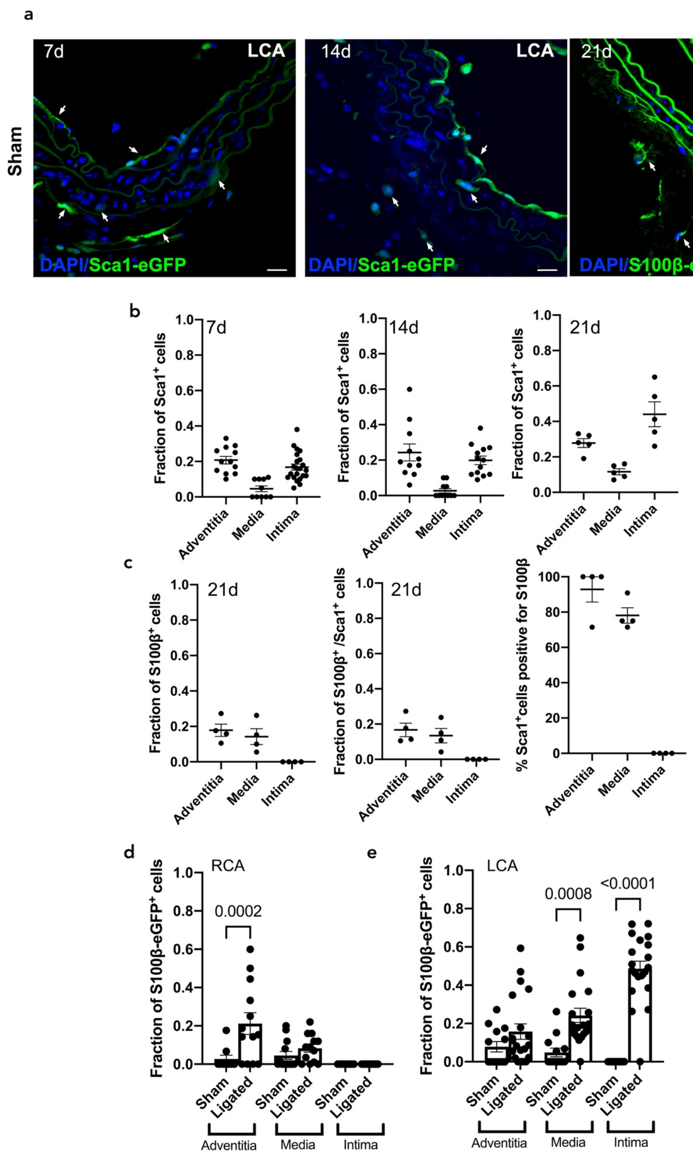


**Suppl Figure 2 (related to Figure 2)**. **Sca1 and S100β cells in adventitial, medial and luminal layers of normal murine arteries**.

**a.** Representative confocal images of Sca1+-eGFP cells in the adventitial, medial and intimal layer of normal sham vessels after 7 days (right panel) and 14 days (middle panel) and S100β-eGFP+ cells after 21 days (right panel). **b.** Cumulative analysis of the fraction of Sca1+ cells in the adventitial, medial and intimal layer of sham vessels after 7, 14 and 21 days. Data are the mean ± SEM, n=5-15 sections from 3 animals/group. **c.** Cumulative analysis of the fraction of S100β+ cells, the fraction of S100β/Sca1+ cells and the percentage of Sca1 cells positive for S100β in the adventitial, medial and intimal layer of sham vessels after 21 days. Data are the mean ± SEM, n= 3 animals/group. Cumulative analysis of the fraction of S100β-eGFP+ cells in the adventitial, medial and intimal layer of **d.** RCA and **e.** LCA vessels, respectively, after 21 days in S100β-eGFP mice. Data are the mean ± SEM, n= 15-20 sections from 3-6 animals/group.

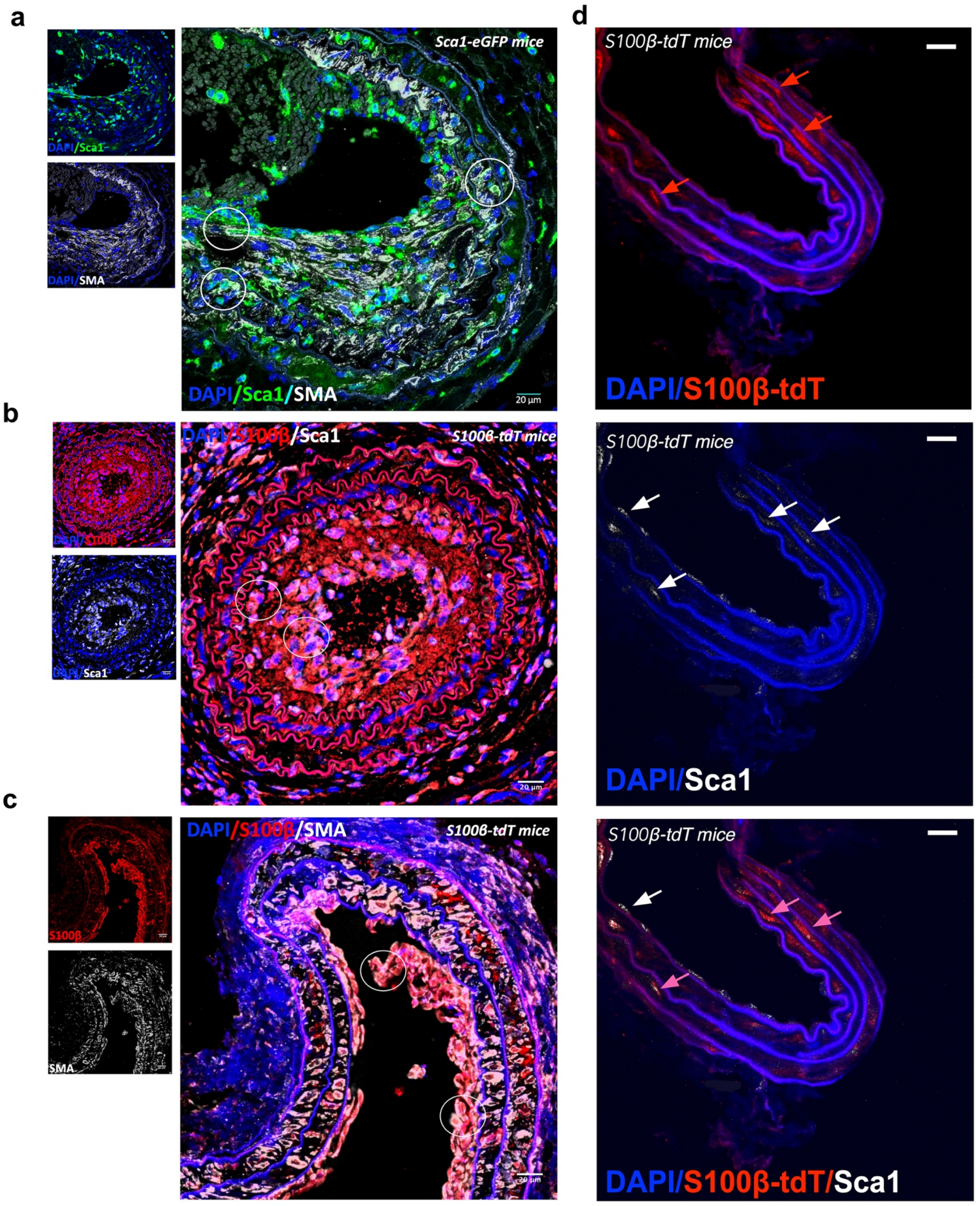


**Suppl Figure 3 (related to Figure 3). Co-localisation of Sca1, S100β and SMA cells in adventitial, medial and luminal layers of injured murine arteries.**

**a.** Representative images of Sca1 (green, left top panel) and SMA (white, left bottom panel) and double stained Sca1/SMA cells (right panel) in ligated LCA of Sca1-eGFP mice after 14 days. **b.** Representative image of S100β-tdT (red, left top panel) and Sca1 (white, left bottom panel) and double stained S100β/Sca1 cells (right panel) of ligated LCA in S100β-CreERT2-tdT mice after 21 days. **c.** Representative image of S100β-tdT (red, left top panel) and SMA (left bottom panel) and double stained S100β/SMA cells (right panel) of ligated LCA in S100β-CreERT2-tdT mice after 21 days**.** Circles are representative of co-localisation. **d.** Representative images of S100β-tdT (red, left panel), Sca1-Far Red (white, middle panel) and double stained S100β/Sca1 (right panel) in the contralateral RCA after 21 days. Data are representative of 6 animals/group.

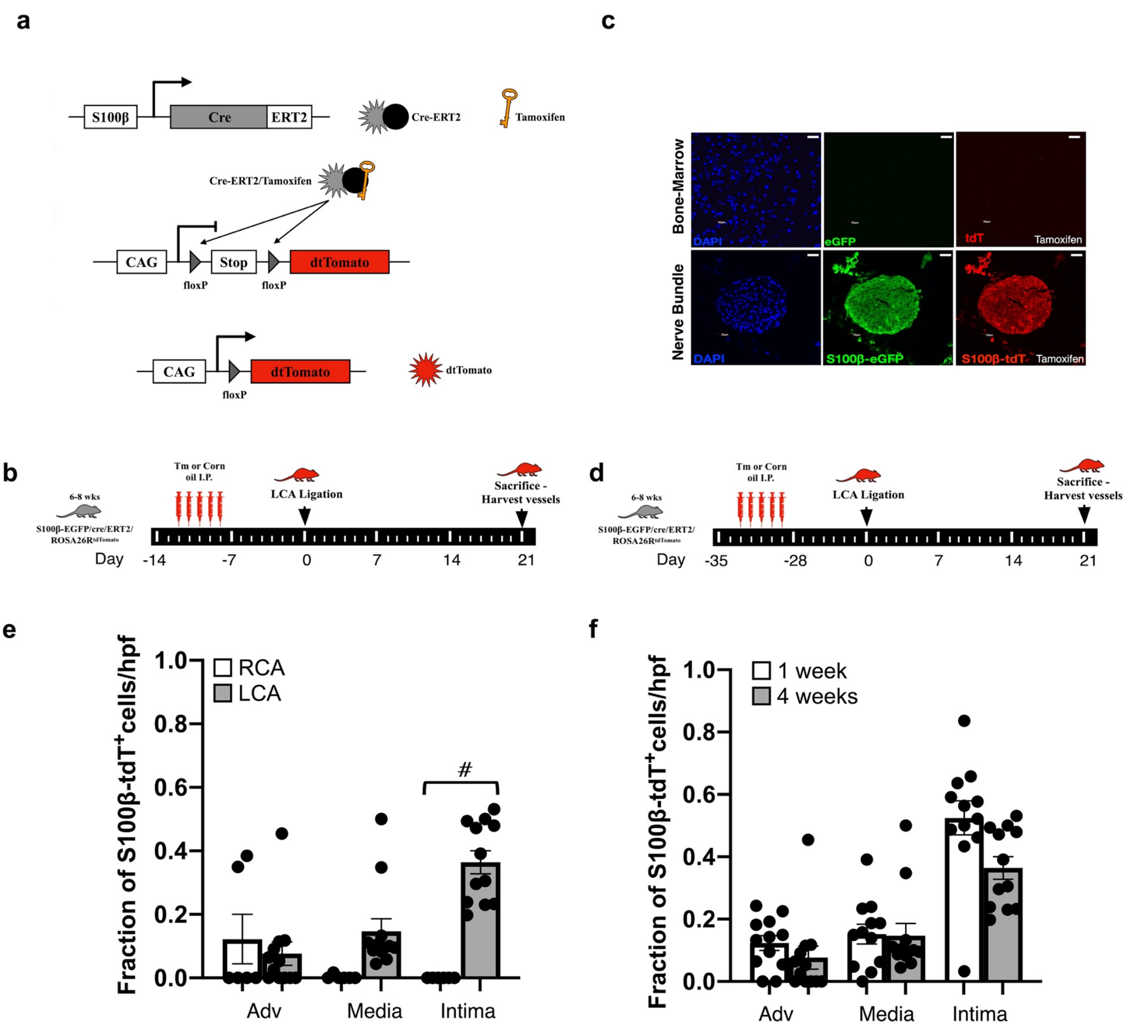


**Suppl Figure 4 (related to Figure 3). Lineage tracing analysis of S100β stem cells**

**a.** Schematic diagram showing generation of S100β-CreERT2-Rosa26-tdTomato. **b**. Tm protocol of 1 week washout prior to ligation. **c.** Representative confocal fluorescence images of S100β-tdT expression in bone marrow smears and herrings nerve bundle in S100β-CreERT2-Rosa26-tdTomato **d.** Tm protocol of 4 week washout prior to ligation. **e.** The fraction of S100β-tdT cells within the adventitial, medial, intimal layers of LCA compared to RCA after 4 week washout. **f.** Comparison of the fraction of S100β-tdT cells within the adventitial, medial, intimal layers of LCA between 1 week and 4 week Tm washout protocols. Data are the mean ± SEM of 3-5 representative images from 3 animals, #$p<0.05$ vs RCA controls.

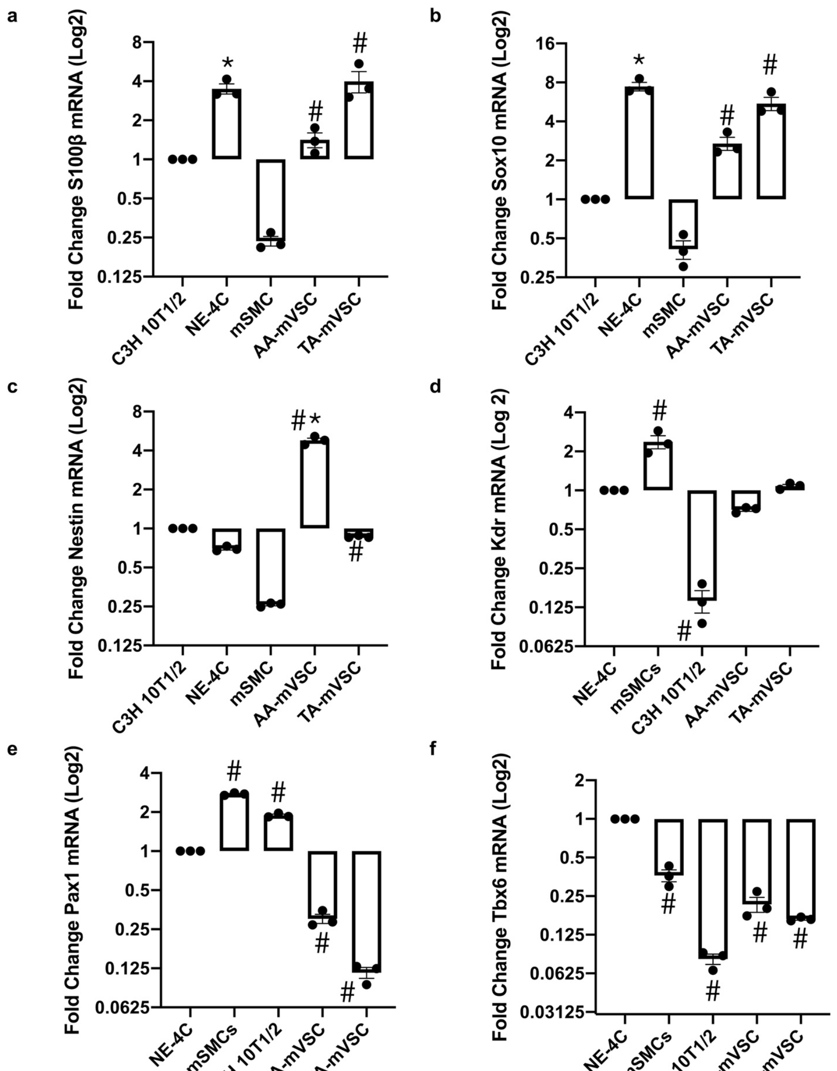


**Suppl Figure 5 (related to Figure 5). Neuroectoderm and paraxial mesoderm marker expression in murine vSCs *in vitro***

**a-c.** Relative levels of neuroectoderm markers (S100β, Sox10 and Nestin) and **d-f.** paraxial mesoderm markers (Kdr, Pax1 and Tbx6) in vSCs isolated from AA and TA regions of the mouse aorta. Data are expressed as the Log2 fold change in mRNA levels relative to C3H 10T1/2 cells or neural stem cells (NE-4C) in culture and are the mean ± SEM of 3 independent cultures, *p≤ 0.05 vs C3H 10T1/2 cells (a-c), #p≤ 0.05 vs NE-4C (d-f).

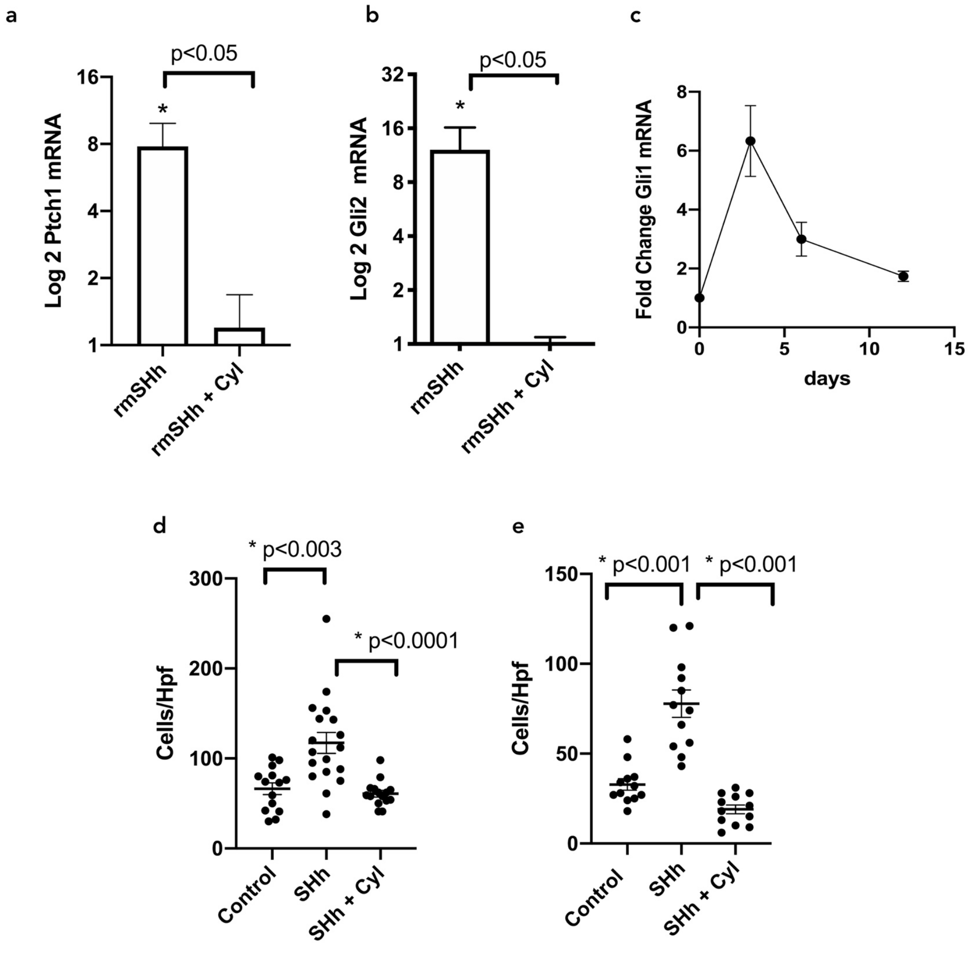


**Suppl Figure 6 (related to Figure 4 and 6). Hedgehog target gene expression and growth response to recombinant SHh.**

Relative levels of **a.** Ptch1 and **b.** Gli2 in vSCs in the presence of rSHh (0.5 µg/ml) with or without the smoothened inhibitor, cyclopamine (10µM). Data are expressed as the Log2 fold change in mRNA levels relative to vSCs alone (control) and are the mean ± SEM of three representative wells, *p≤0.05 versus control. **c.** Temporal fold increase in Gli1 mRNA levels following carotid ligation. Data are the mean ± SEM from 3 vessels. The effect of recombinant SHh on the growth of **d.** S100β$^{+}$ vSC and **e.** S100β$^{+}$ NEPs in culture after 7 and 12 days, respectively. Data are the mean ± SEM of 8-12 wells/group.
